# SFUMATO: Bayesian probabilistic clustering for uncertainty-aware spatial transcriptomics analysis and mapping

**DOI:** 10.64898/2026.09.25.754409

**Authors:** Alessia Giustolisi, Christophe Avenel, Carolina Wählby

## Abstract

In situ sequencing methods provide subcellular-resolution gene expression data while preserving spatial context, enabling the integrated analysis of genomics and tissue morphology. However, many clustering workflows represent spatial transcriptomic structure through hard labels, imposing sharp boundaries between transcriptional domains and limiting the visualisation of gradual transitions, mixed signals, and uncertainty in cluster assignment, resulting in potentially misleading interpretations of tissue organisation.

Here we present SFUMATO (Segmentation-Free Uncertainty Mapping and Analysis of Transcriptomic Organisation), a GPU-accelerated Bayesian probabilistic clustering and visualisation framework for spatial transcriptomics. SFUMATO builds on segmentation-free, multi-scale transcript binning and applies a Bayesian Gaussian mixture model to infer posterior probabilities over transcriptional components at each spatial location. These posterior probabilities provide a unified representation for hierarchical clustering, uncertainty-aware visualisation, and quantitative estimation of transcript-defined area fractions.

SFUMATO assigns related colours to transcriptionally related components and blends colours according to posterior probabilities, producing maps in which sharp boundaries, gradual transitions, and ambiguous regions are represented directly in the visualisation. Classical hard labels remain recoverable from the same posterior distributions, preserving compatibility with downstream analyses that require discrete clusters.

We evaluate SFUMATO on public 10x Genomics Xenium mouse brain and human breast cancer datasets. In mouse brain, SFUMATO preserves local transcriptional similarity in its colour encoding, maintains hierarchically consistent visualisations across different numbers of clusters, and recovers cell-associated spatial organisation without requiring prior cell segmentation. In breast cancer, SFUMATO-derived posterior maps can be converted into semantic masks concordant with pathology-associated regions, particularly invasive carcinoma, while highlighting the greater heterogeneity of DCIS-associated tissue.

Together, these results show that SFUMATO provides an interpretable probabilistic representation of spatial transcriptomics that connects clustering, visualisation, and semantic segmentation across cell-associated and niche-level scales.

More broadly, SFUMATO also provides a compact and continuous representation of highly multiplexed spatial transcriptomic information, offering a flexible input for future downstream analyses and multimodal integration.

## 1 Introduction

In the context of spatial omics, spatial transcriptomics technologies enable the visualisation and analysis of gene expression in conjunction with tissue imaging. Among these approaches, *in situ* sequencing (ISS) provides subcellular-resolution, while preserving spatial information and achieving a high degree of transcript multiplexing.

Recent advances, such as Xenium, generate datasets comprising millions of spatially resolved transcripts across hundreds to thousands of genes, representing gene expression as a dense point process in a pseudo-continuous space. Following acquisition and decoding, these data are typically represented as lists of decoded transcripts with coordinates mapping each transcript to a spatial position in the imaging domain, thereby enabling highly multiplexed spatial analyses [1].

To explore latent structures within such data, which are inherently high-dimensional and difficult to interpret through visual inspection alone, classical clustering techniques and their more advanced variants are commonly employed. These methods partition the gene expression space into discrete regions, which are subsequently projected onto the spatial domain for visualisation. However, biological patterns are often characterised by gradual transitions, poorly represented by sharp-edged regions. Moreover, “hard” clustering frequently introduces boundary effects at the interfaces between spatial domains.

To address these limitations from a visual perspective, Partel *et al*. [2] proposed a colour-based encoding of gene expression variation, where reduced feature representations obtained through dimensionality reduction techniques such as Uniform Manifold Approximation and Projection (UMAP) [3] are mapped to RGB colour space and projected back onto the spatial domain. This approach provides an intuitive visualisation of gradual transitions, highlighting spatial continuity and regional variations in gene expression patterns, but does not result in a partitioning of the tissue space that can be used for downstream processing.

Complementary to visual strategies, probabilistic models have been proposed to capture soft structure in spatial transcriptomics data more rigorously. FICTURE [4] is a segmentation-free method that models spatial gene expression using a probabilistic framework in which each location is represented as a mixture of latent transcriptional programs. While these mixtures are often visualised by assigning each location to its dominant factor, yielding cluster-like maps, the inferred factors are not explicitly defined as spatial clusters and require post hoc interpretation. Bayesian approaches such as BFAST [5] offer rich statistical modelling through joint dimensionality reduction and clustering via factor analysis, accounting for complex covariance structures and technical effects such as zero inflation. However, the computational burden of such modelling strategies limits their application to datasets comprising tens of millions of spatially resolved transcripts.

One approach to reduce both data dimensionality and the number of data points is to first try to identify individual cells. This, in turn, raises the challenge of cell segmentation, often relying on cell nuclei that may be poorly defined in, e.g., cancer tissue. GraphST [6] is an example of a segmentation-based approach, as it operates on cell-level representations. Segmentation-free approaches bypass this requirement by operating directly on the spatial distribution of transcripts, thereby reducing dependence on the errors and assumptions introduced by cell boundary delineation. On the other hand, the output from a segmentation-free approach will not directly reflect the number of data points.

To enable quantitative analysis of spatial transcriptomics data represented as point processes, feature extraction is required to transform individual transcript coordinates into a structured representation of local gene expression. Specifically, the spatial domain is converted into a feature matrix in which each observation encodes the gene composition within a defined local spatial neighbourhood. Several methods have been developed, including probabilistic factorisation [4, 7], graph-based spatial clustering [8], and bin-based aggregation strategies [9]. Among these, Points2Regions [9] is particularly relevant because it provides a fast and scalable segmentation-free representation of transcript-level data. It aggregates transcripts into spatial bins, applies Gaussian smoothing, and performs k-means clustering followed by hierarchical clustering on normalised multiple-resolution bin composition vectors. As demonstrated by Salas *et al*. [10] on Xenium mouse brain data, Points2Regions is well suited to high-transcript-density datasets and can reveal molecular signatures at both cell and tissue-niche scale without relying on cell boundary delineation.

Its scalability and bin-based formulation make Points2Regions particularly useful for rapid parameter exploration and interactive inspection of spatial transcriptomic structure, including through visualisation environments such as TissUUmaps [11, 12]. However, as a hard clustering method, it assigns each bin to a single cluster, precluding direct representation of uncertainty or gradual transitions between transcriptional domains.

From a quantitative perspective and for further analysis of tissue composition, a deeper limitation of segmentation-based approaches lies in what their outputs actually measure. Cell-type counts produced by segmentation are not necessarily proportional to the underlying volume fractions of the same cell types in tissue, making comparisons to cell counts from single cell sequencing skewed. By contrast, the Delesse principle, in its original formulation, states that the areal proportion of a given phase observed on a planar section is an unbiased estimator of its volumetric proportion in the bulk material [13]; through domain shift, it can be claimed that area fractions measured in a histological section are proportional to volume fractions in the underlying three-dimensional structure. This suggests that area-based estimates of cell-type composition, rather than cell counts, provide a more principled approximation of volumetric composition. A probabilistic, segmentation-free representation of spatial gene expression – in which each spatial location carries a soft assignment over transcriptional clusters – is naturally interpretable as an estimate of local area fractions, and therefore provides a more direct path to Delessebased compositional inference than hard segmentation or hard clustering. We acknowledge that this correspondence is subject to geometric and sampling biases when applied to biological tissues [14].

Here we present SFUMATO (Segmentation-Free Uncertainty Mapping and Analysis of Transcriptomic Organisation), a GPU-accelerated Bayesian probabilistic clustering and visualisation framework designed to represent the highly multiplexed functional information of spatial transcriptomics datasets in intuitive maps of biological complexity. By exploiting posterior probabilities, SFUMATO generates transcriptomic maps in which similar colours indicate similar gene-expression composition, while colour mixing reflects uncertainty in cluster assignment. This allows transition zones and mixed transcriptional states to be visualised without false and unstable partitioning from hard clustering.

The name SFUMATO is inspired by the Renaissance painting technique, most notably associated with Leonardo da Vinci, in which subtle changes of tone are used to soften transitions and avoid sharp boundaries. In an analogous manner, SFUMATO replaces hard cluster assignments with posterior probability distributions over latent transcriptional components, allowing spatial domains to be represented as continuous and potentially overlapping. By encoding both similarity in gene expression and uncertainty in cluster assignment, SFUMATO aims to produce representations in which related biological structures appear visually coherent, while transitional zones are explicitly revealed rather than suppressed.

The main contributions of SFUMATO are as follows:

1. **Segmentation-free visualisation of local transcriptional structure**. SFUMATO provides an efficient representation of transcript-level spatial data without requiring prior cell segmentation. By linking colour assignment to the expression-based hierarchy of cluster composition, SFUMATO produces visual maps in which colour similarity reflects transcriptional similarity. We show that this colour encoding captures local gene-expression similarities and differences at cell-associated scale, in comparison with methods such as Points2Regions and FICTURE.
2. **Hierarchically consistent visualisation across clustering resolutions**. SFUMATO derives multiple clustering resolutions from a shared overclustered probabilistic model and a common expression-based dendrogram. As a result, changing the number of clusters *K* refines the same hierarchical representation rather than producing unrelated categorical maps. This enables stable visualisation of both gradual transitions and sharp transcriptional boundaries coherently across different values of *K*.
3. **Probabilistic clustering for semantic segmentation across biological scales**. SFUMATO retains posterior probabilities over transcriptional components and uses them to derive semantic regions from transcript-defined patterns. At cell-associated scale, these regions can be compared with segmentation-derived cell-type annotations while remaining independent of the cell segmentation itself. At niche scale, posterior probability maps can be converted into semantic masks and compared with pathology-derived tissue annotations, supporting transcript-driven segmentation of pathology-associated regions. Because SFUMATO models the spatial distribution of transcripts directly rather than aggregating them within predefined cell boundaries, it may also reveal subcellular or pericellular transcriptional patterns that are hidden by methods relying on cell segmentation.

Building on the Points2Regions feature extraction pipeline, SFUMATO applies a Bayesian Gaussian Mixture model (BGM) to a low-dimensional representation of binned transcript profiles. The resulting posterior probabilities provide a unified representation that supports multiple downstream uses:

- **Uncertainty-aware clustering**. Each spatial bin is assigned a full posterior distribution over *K* clusters rather than a single hard label, allowing uncertainty to be quantified at domain boundaries and in regions of mixed cellular composition.
- **Hierarchical visualisation**. BGM components are organised according to the expression-based hierarchy of their posterior-weighted soft centroids. Cluster colours are assigned from this hierarchy, so clusters with related transcriptional components receive related colours, while distant components are shown with stronger colour contrast.
- **Probabilistic colour mixing**. For visualisation, the colours of the two most probable components are blended according to their posterior probabilities. Sharp boundaries emerge where the posterior mass is concentrated on a single component, whereas gradual colour transitions reveal regions of biological ambiguity or mixed transcriptional signal.
- **Semantic segmentation and area quantification**. Posterior probability maps can be converted into semantic masks and area fractions, providing quantitative estimates of the spatial extent of transcript-defined tissue components.
- **Compatibility with hard-label workflows**. Classical hard cluster labels remain recoverable from the same posterior distributions by assigning each bin to its maximum-posterior component, preserving compatibility with downstream pipelines that require discrete labels.

We evaluate SFUMATO against methods representing complementary analysis assumptions: GraphST and the Xenium 10x Genomics workflow as segmentation-based cell-level references, FICTURE as a segmentation-free probabilistic factorisation framework, and Points2Regions as the closest bin-based hard-clustering baseline. Together, these comparisons assess whether SFUMATO preserves transcriptomic similarity in its colour encoding, maintains hierarchical consistency across clustering resolutions, and supports semantic segmentation at both cell-associated and niche scales.

## 2 Methods

The SFUMATO workflow consists of feature extraction and preprocessing, BGM over-clustering, dendrogram cut, colour assignment, and probabilistic colour mixing, as illustrated in Figure 1 and detailed below.

**Figure 1:**
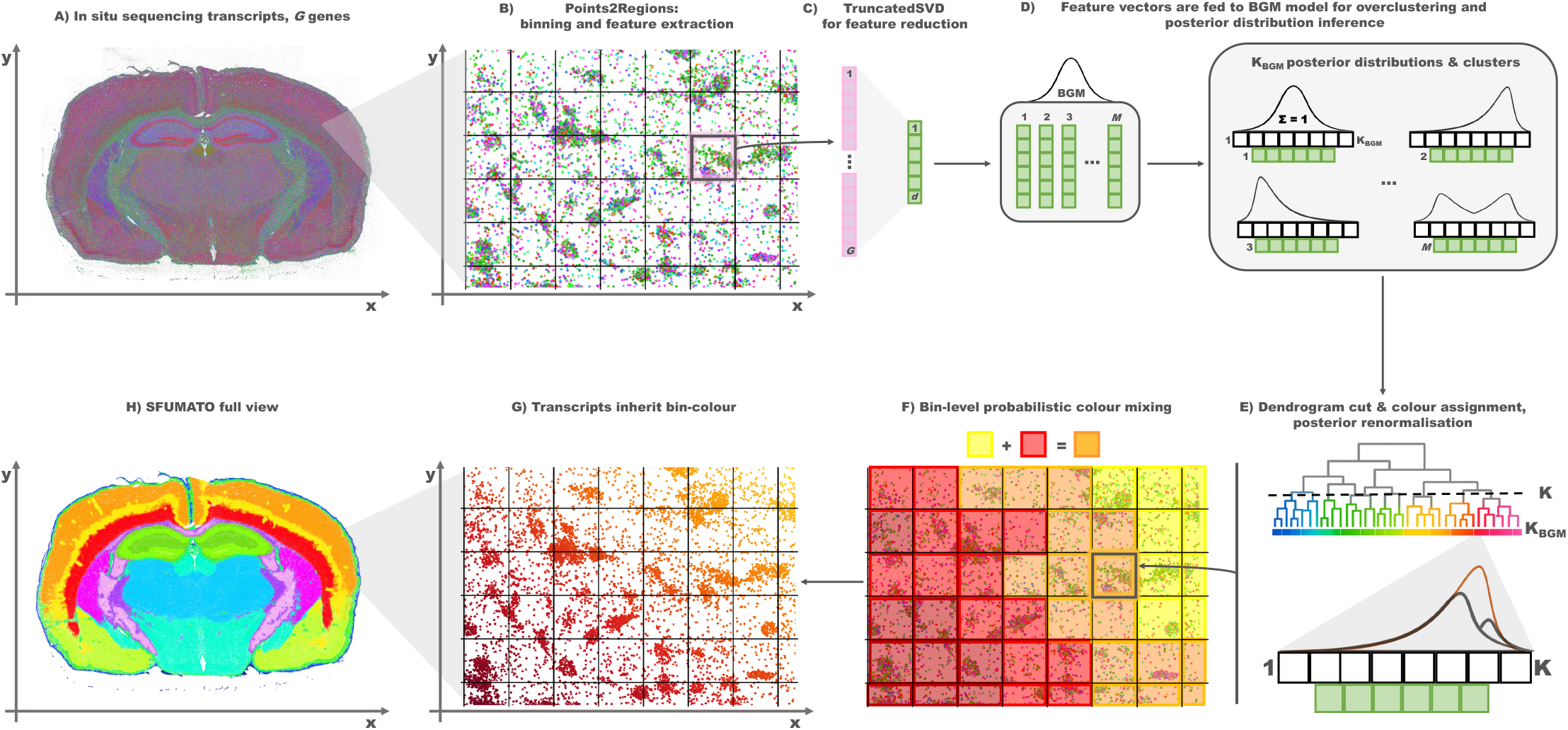
Overview of the SFUMATO workflow. (**A**) In situ sequencing data are represented as *G* detected genes with spatial coordinates in the image plane. (**B**) A Points2Regions-inspired binning and feature extraction step converts transcript coordinates into local gene-expression feature vectors. (**C**) The resulting sparse feature matrix is reduced to *d* latent components using TruncatedSVD. (**D**) The reduced feature vectors are used as input to an overclustered BGM, which infers posterior cluster probabilities for each spatial bin. (**E**) Overclustered components are organised in an optimally ordered dendrogram; cutting this dendrogram yields the requested final clusters, posterior probabilities are renormalised after merging, and cluster colours are assigned according to the dendrogram ordering. (**F**) At bin level, posterior probabilities are used for logarithmic probabilistic colour mixing. (**G**,**H**) For visualisation, transcripts inherit the colour assigned to their corresponding bin.

### 2.1 Feature extraction and preprocessing

We adopt a Points2Regions-like feature extraction pipeline [9] (Fig. 1B), a computationally efficient and segmentation-free approach designed for large-scale spatial transcriptomics datasets. Transcripts are rasterised onto square grids at multiple spatial resolutions, producing class-count representations of gene expression. Each transcript is assigned to a spatial bin via a sparse binning operator, and gene identities are encoded through one-hot representations.

Feature vectors are extracted at locations corresponding to the centres of the highest-resolution grid. To integrate information across spatial scales and mitigate discretisation artefacts, class counts from neighbouring pixels at different resolutions are aggregated using inverse-distance weighted interpolation. This yields a multi-scale representation of local gene composition, analogous to an image pyramid, capturing both fine-grained and coarse spatial structure.

Following feature extraction, bins with insufficient transcript multi-scale density are filtered out, yielding a set of valid spatial locations (hereafter referred to as *good bins*). Let *M* denote the number of good bins and *G* the number of genes. The resulting feature matrix is denoted as *X* ∈ R^*M* ×*G*^, where each row vector **x**_*i*_ represents the local gene expression composition of the good bin *i*.

To stabilise the representation and account for variability in transcript counts, the smoothed binned gene counts are log-transformed using a log(1+*x*) mapping and subsequently L1-normalised across genes within each bin, yielding compositional feature vectors that are comparable across spatial locations.

Given the high dimensionality and sparsity of the gene space, dimensionality reduction is performed using Truncated Singular Value Decomposition (TruncatedSVD) [15, 16] (Fig. 1C). Unlike conventional Principal Component Analysis (PCA), TruncatedSVD operates directly on sparse matrices without requiring matrix densification, making it particularly suitable for large, highly multiplexed spatial transcriptomics datasets. The transformation is learned from the data and applied to obtain a low-dimensional latent representation *Z* ∈ R^*M* ×*d*^, where *d* denotes the number of retained latent components. The parameter *d* is user-configurable and can be adapted to the complexity of the dataset and the desired level of feature compression. Each vector **z**_*i*_ corresponds to the projection of **x**_*i*_ onto the resulting latent embedding.

### 2.2 Bayesian Gaussian Mixture Model

Clustering is formulated as variational Bayesian inference in a Bayesian Gaussian Mixture (BGM) model [17] applied to the latent representation obtained by TruncatedSVD (Fig. 1D). BGM introduces prior distributions over all parameters to address the limitations and over-fitting issues associated with standard Gaussian Mixture Models. Let *Z* ∈ R^*M* ×*d*^ denote the resulting latent feature matrix, where *M* is the number of spatial bins, *d* is the number of retained latent dimensions, and each row **z**_*i*_ ∈ R^*d*^ represents the low-dimensional embedding of spatial bin *i*. Given a target number of mixture components *K*_BGM_, each spatial bin is assumed to arise from one of *K*_BGM_ latent transcriptional components,

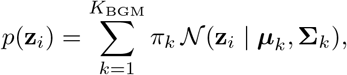

where *π*_*k*_ denotes the mixture weight of component *k*, ***µ***_*k*_ is the mean vector of the corresponding multivariate Gaussian distribution, and **Σ**_*k*_ is its covariance matrix.

The model was implemented following the variational Bayesian formulation used by the `BayesianGaussianMixture` class in `scikit-learn` [16], but re-implemented in PyTorch to enable GPU execution. In practice, this BGM avoids Markov chain Monte Carlo sampling and instead approximates posterior distributions through variational inference, making the approach computationally tractable for large datasets containing millions of spatial bins, particularly at small bin widths used for cell-scale analysis.

For computational efficiency, diagonal covariance matrices were used:

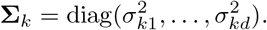

This approximation assumes conditional independence between latent dimensions within each component, thereby de-coupling covariance estimation across dimensions. Although less flexible than full covariance modelling, the diagonal formulation substantially reduces memory usage and computational complexity, while preserving sufficient flexibility in the embedded latent space.

Variational inference proceeds through an expectation– maximisation (EM)-like iterative procedure. During the responsibility update, the posterior responsibility of each mixture component is computed for every spatial bin,

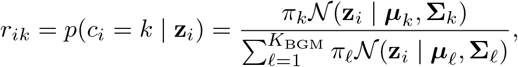

where *c*_*i*_ denotes the latent component assignment of spatial bin *i*, and *r*_*ik*_ is the posterior probability that bin *i* belongs to component *k*. These posterior probabilities, commonly referred to as *responsibilities*, quantify the contribution of each mixture component to the observed latent embedding **z**_*i*_.

The expression above provides the standard Gaussian-mixture interpretation of the responsibilities. In the Bayesian Gaussian mixture used here, the mixture weights and Gaussian parameters are themselves described by variational posterior distributions rather than treated as fixed point estimates. Accordingly, the responsibility updates are based on the corresponding expected log mixture weights and expected Gaussian log-likelihoods under these variational distributions.

The resulting responsibilities are then used to update the variational posterior distributions over the mixture weights, component means, and covariances, namely {*π*_*k*_, ***µ***_*k*_, **Σ**_*k*_}, rather than point estimates of these quantities. These variational updates were performed in single precision (float32) on GPU.

Model fitting was initialised using k-means++ [18] unless otherwise specified. The variational inference procedure was iterated until convergence of the evidence lower bound (ELBO), stopping when the absolute change in lower bound between consecutive iterations fell below a tolerance threshold. A maximum number of iterations was set, but convergence was always reached before this limit.

The primary output of the model is the posterior responsibility matrix

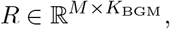

where each entry *r*_*ik*_ represents the posterior probability that bin *i* belongs to component *k*.

Consistent with the probabilistic formulation, SFUMATO represents each mixture component by a soft centroid computed from the posterior responsibilities:

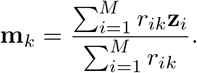

This definition allows each bin to contribute to multiple centroids proportionally to its posterior assignment probability, rather than being assigned exclusively to a single cluster. Unlike the Gaussian means ***µ***_*k*_, which are internal parameters of the Bayesian mixture model, these posterior-weighted centroids summarise the observed data associated with each component and are subsequently used to quantify similarities between components during the hierarchical merging step.

Hard cluster labels can be recovered as

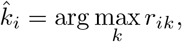

while the posterior probabilities remain available for uncertainty-aware visualisation and other downstream probabilistic analyses.

### 2.3 Merging and colour assignment

Although a BGM can be used as a standalone clustering algorithm to obtain a single clustering with a user-defined number of components, SFUMATO’s main strength lies in deriving multiple, hierarchically consistent clusterings from a single probabilistic model (Fig. 1E).

The BGM employs diagonal covariance matrices, assuming conditional independence between latent dimensions. While this approximation provides substantial computational gains, it may limit the ability of the model to capture more complex cluster geometries. Following the rationale of Points2Regions [9], SFUMATO therefore adopts an overclustering strategy, in which a larger number of mixture components is first estimated and subsequently merged according to their transcriptional similarity.

Specifically, when the user requests one or more clusterings *K*_1_, …, *K*_*n*_, the BGM is fitted only once using

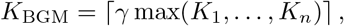

where *γ >* 1 is an oversampling factor. Throughout this work, we use *γ* = 1.5, following the empirical observations reported by Andersson et al. [9].

Unlike conventional BGM, where components with negligible mixture weights may be removed through pruning, SFUMATO retains all overclustered components. The resulting overclustered model defines a shared hierarchical representation from which all requested clusterings are subsequently obtained.

After fitting the BGM, pairwise cosine distances are computed between the soft centroids of the overclustered components, and an agglomerative hierarchical clustering is performed using complete linkage with optimal leaf ordering. The latter produces a one-dimensional ordering of the leaves that is consistent with the hierarchical structure while minimizing the distances between adjacent leaves, providing a natural ordering for subsequent colour assignment. The shared dendrogram is then cut at each requested value of *K*, yielding the corresponding merged clustering without requiring additional BGM fits (Fig. 1E). The merged posterior probabilities are finally renormalised so that they again form a valid probability distribution.

The motivation for this strategy is twofold. First, deriving all requested clusterings from the same probabilistic model ensures that relationships between clusters remain hierarchically consistent across different values of *K*. Second, the same shared hierarchy forms the basis of the colour assignment strategy, linking the hierarchical organisation of the clusters directly to their visual representation.

Each merged cluster occupies a contiguous span of leaves along the optimally ordered dendrogram. A representative position is assigned to each cluster as the centre of this span and mapped onto a continuous colour map. Consequently, neighbouring colours are assigned to clusters that are nearby in the expression-based hierarchy, whereas increasingly distinct colours correspond to progressively more distant branches of the hierarchy. Because every requested clustering is obtained by cutting the same optimally ordered dendrogram, colour assignment remains hierarchically consistent across different values of *K*.

By default, colours are sampled from the `gist ncar` colourmap after trimming its extreme endpoints to improve visual separability. Users may alternatively select continuous colourmaps, including perceptually uniform, colour-vision-deficiency-friendly options such as `viridis` and `cividis` [19], without affecting the underlying clustering procedure.

### 2.4 Logarithmic colour mixing

The probabilistic formulation of SFUMATO is carried through to the final visualisation. Posterior probabilities estimated at the spatial-bin level are used to assign colours according to each bin’s posterior distribution (Fig. 1F), rather than the single most probable cluster, thereby encoding uncertainty in the visual representation (Fig. 1G and H).

Let *r*_*i*1_ and *r*_*i*2_ denote the two highest posterior probabilities for spatial bin *i*, corresponding to clusters with colours **c**_1_ and **c**_2_ assigned through the hierarchical colour assignment described above. The final colour **c**_*i*_ assigned to bin *i* is defined as

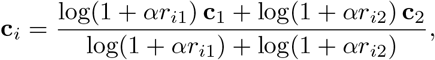

with *α* = 50.

Colour mixing is performed directly on the RGB coordinates returned by the selected colourmap, to avoid artefacts that may arise when interpolating colours directly in HSV space because of the circular nature of the hue component. The logarithmic weighting increases the visual contribution of secondary cluster memberships while preserving the dominance of high-confidence assignments. For visualisation, all transcripts belonging to the same spatial bin inherit the colour assigned to that bin (Fig. 1G and H). Transcripts belonging to bins with insufficient transcript density to be involved in the clustering are assigned a neutral black colour.

As a result, transcripts belonging to spatial bins with ambiguous cluster membership, particularly at transition zones, are represented by smooth colour mixtures, whereas well-separated transcriptional domains retain distinct, un-mixed colours. Combined with the hierarchical colour assignment, this probabilistic colour mixing provides a visual representation in which both transcriptional similarity and uncertainty are directly encoded.

### 2.5 Data and method evaluation

We evaluated SFUMATO on two publicly available Xenium ISS transcriptomics datasets: a fresh frozen mouse brain section profiled with the Xenium Mouse Brain Gene Expression Panel, comprising more than 40 million detected transcripts across 248 genes [20], and a human breast cancer formalin-fixed, paraffin-embedded (FFPE) section from the Xenium human breast cancer dataset, profiled with the 280-gene Xenium Human Breast Gene Expression Panel and 33 additional custom genes, for a total of 313 targeted genes and a total of 42, 433, 535 transcripts [21, 22]. Dataset details are reported in Supplementary Table 1.

For comparisons between SFUMATO and other clustering methods at the cell level, we used row-normalised transcript-level co-occurrence matrices. Since SFUMATO operates at the bin level, each transcript was assigned the maximum-posterior cluster of its corresponding bin. For each comparison, matrix entries were computed by counting the number of transcripts assigned to each pair of SFUMATO and reference labels. Rows were normalised to sum to one, so that each entry represents the fraction of transcripts from a given SFUMATO cluster assigned to a given cluster or factor by the comparison method.

To compare the probabilistic output of SFUMATO with reference spatial annotations and segmentations, we first converted the posterior probabilities into binary semantic segmentation masks. For each cluster, the full posterior distribution was projected onto the spatial domain by assigning to every location within a spatial bin the posterior probability of that bin, yielding a bin-level posterior probability map for each cluster. The probability maps were subsequently smoothed with a Gaussian filter whose characteristic scale was chosen to match the spatial bin size.

Binary semantic regions were then extracted using a two-threshold morphological reconstruction procedure. Pixels with probabilities above a high threshold were first selected as confident seeds, while pixels above a lower threshold were retained only if connected to one of these seeds through morphological reconstruction. This procedure suppresses isolated low-probability regions while preserving spatially coherent semantic domains.

Agreement between SFUMATO-derived semantic regions (*A*) and reference masks (*B*) was quantified using the Dice similarity coefficient,

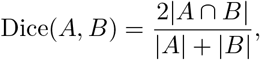

where |.| denotes the number of pixels in the corresponding binary masks. This analysis evaluates the spatial overlap between semantic regions derived from the probabilistic clustering rather than direct correspondence between individual transcript labels.

For comparisons with the Xenium analysis pipeline, reference masks were generated from the cell segmentation and cell type annotations provided by 10x Genomics. For each annotated cell type, all segmented cells belonging to that cell type were merged into a single binary mask, yielding one reference semantic region per cell type. Dice similarity was then computed between these reference masks and the corresponding SFUMATO-derived semantic regions.

## 3 Results

### 3.1 SFUMATO colour encoding reflects local transcriptional similarity

We first evaluated whether SFUMATO preserves transcriptomic similarities and provides an interpretable segmentation-free visualisation of local transcriptional structure. SFUMATO was run on the Xenium mouse brain dataset at a bin width selected to resolve cell-scale expression patterns, with parameters reported in Supplementary Tables 2–3. We compared SFUMATO with two other segmentation-free methods, Points2Regions [9] and FICTURE [4], using the same number of clusters or factors (*K* = 45) – matching the number of clusters proposed by the 10x Genomics analysis workflow for this dataset – and parameters selected to resolve the same scale as SFUMATO. For FICTURE, each anchor point was assigned to the factor with maximum posterior probability, and transcript-level labels were obtained by assigning each transcript to the dominant factor of the corresponding anchor point. Points2Regions was run using the same feature extraction parameters as SFUMATO, while retaining its original k-means clustering followed by hierarchical cutting of the cluster dendrogram.

SFUMATO, Points2Regions, and FICTURE produced broadly concordant spatial organisations at this scale (Fig. 2A). Using the transcript-level co-occurrence analysis defined above, SFUMATO showed substantial agreement with both Points2Regions and FICTURE, while also revealing many-to-one correspondences, particularly in the comparison with FICTURE. These differences are expected given that FICTURE uses a distinct anchoring strategy and a Latent Dirichlet Allocation-based probabilistic factor model, whereas SFUMATO and Points2Regions share the same feature extraction stage. Importantly, many-to-one correspondences often involve SFUMATO clusters with similar colours, indicating that the labels collapsed into a single comparison cluster are neighbours in the SFUMATO expression-based hierarchy and therefore transcriptionally related, rather than arbitrary or unrelated subdivisions.

**Figure 2:**
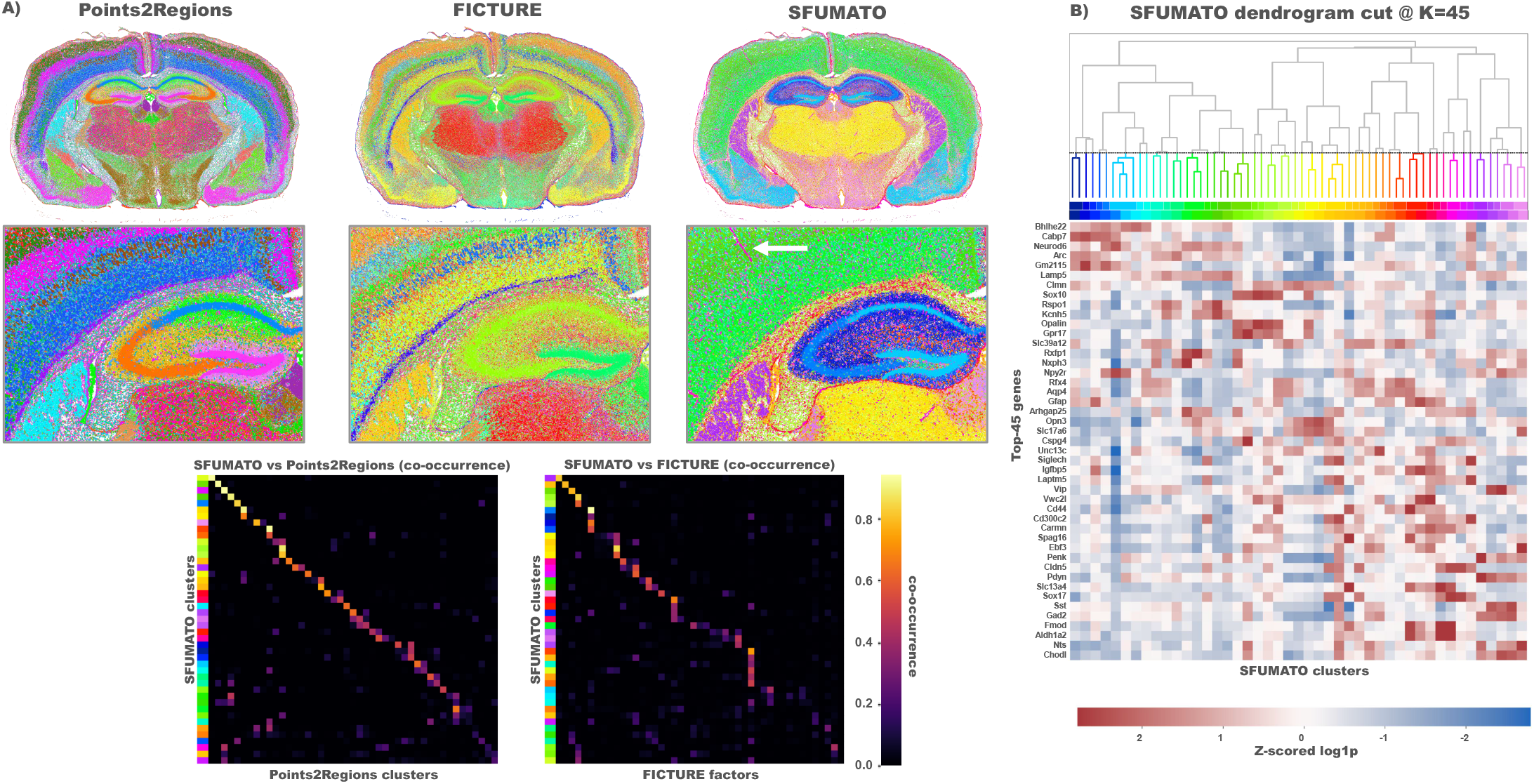
SFUMATO colour encoding reflects local transcriptional similarity. (**A**) SFUMATO colour encoding reflects local transcriptional similarity in the Xenium mouse brain dataset. Full brain and zoomed region are shown for SFUMATO, Points2Regions, and FICTURE at *K* = 45 clusters or factors. Row-normalised transcript-level co-occurrence matrices compare SFUMATO hard assignments with Points2Regions clusters and FICTURE factors. (**B**) The SFUMATO dendrogram shows the shared expression-based hierarchy used for colour assignment, with clusters ordered according to the optimal leaf order. The differential gene expression heatmap shows 45 marker genes across SFUMATO clusters, with cluster columns and the colour strip ordered according to the same dendrogram. Similar SFUMATO colours correspond to neighbouring positions in the expression-based hierarchy and to related marker-expression profiles, whereas transcriptionally distinct regions are shown with contrasting colours, such as the thin red structure by the white arrow in (**A**), corresponding to a vessel. The cell-scale SFUMATO-Points2Regions and FICTURE results can be explored interactively at https://sfumato.serve.scilifelab.se/xenium_mouse_SFUMATO_cell.tmap and https://sfumato.serve.scilifelab.se/xenium_mouse_FICTURE_cell.tmap, respectively.

We next tested how well clusters reflected similarities and differences in gene expression. For each SFUMATO cluster, we aggregated transcript counts by gene and performed a cluster-level differential gene expression (DGE) analysis to identify enriched marker genes. Candidate cluster–gene associations were filtered according to predefined criteria for positive log-fold change against the remaining clusters, cluster specificity, and within-cluster abundance, and then ranked using a composite score incorporating these three properties. The 45 highest-ranking eligible marker genes were selected subject to a cap on the number of genes associated with any single cluster, thereby preventing clusters with many strong candidates from dominating the panel. Z-scored log1p counts per million for the selected genes were then visualised across SFUMATO clusters. Both the cluster columns and the colour strip above the heatmap were ordered according to the dendrogram leaf order used for SFUMATO colour assignment, while genes were ordered by the centre of their positive expression signal along this axis.

The resulting heatmap (Fig. 2B) shows a structured, approximately diagonal organisation, indicating that clusters with neighbouring SFUMATO colours also tend to share related marker-expression profiles. Conversely, clusters with distinct colours show more divergent marker profiles. This supports the intended role of the SFUMATO colour encoding: similar colours indicate proximity in the shared expression-based hierarchy, whereas strong colour contrasts highlight transcriptionally distinct regions. In the spatial map, this behaviour is visible in regions with deviating gene expression, such as vessel-associated structures, which appeared in red/pink and were clearly contrasted from green-shaded neighbouring clusters (Fig. 2A, white arrow). At the same time, neighbouring green shades reveal gradual differences within the surrounding tissue, illustrating how the SFUMATO colour encoding can display related but non-identical transcriptional states without imposing visually abrupt categorical boundaries. In contrast, purely categorical colour assignments can make such relationships harder to perceive, or can visually exaggerate boundaries between clusters that are transcriptionally related.

### 3.2 SFUMATO preserves gradual and sharp transitions across cluster resolutions

We next evaluated whether SFUMATO provides a stable visual interpretation when the number of clusters is varied. SFUMATO was run on the Xenium mouse brain dataset at a niche-level, so a larger bin width was selected to capture broader tissue-level patterns, with parameters reported in Supplementary Tables 2–3. A single oversampled BGM was fitted using 1.5 × *K*_max_ components, where *K*_max_ = 30, and the shared component dendrogram was cut at *K* = 15, 20, 25, and 30. For the same values of *K*, we ran Points2Regions using the same niche-level feature extraction parameters. This comparison isolates the effect of the probabilistic hierarchical clustering and colour assignment, while keeping the upstream representation comparable.

Across increasing values of *K*, SFUMATO maintained a coherent visual organisation of the mouse brain (Fig. 3), demonstrating that the clustering is hierarchically consistent. This behaviour follows directly from the shared dendrogram: increasing *K* refines existing branches rather than producing an unrelated clustering and colour assignment at each cluster resolution. As a result, patterns that are stable across cuts retain similar colours, while newly separated subclusters appear as related shades when they originate from nearby positions in the expression-based hierarchy.

**Figure 3:**
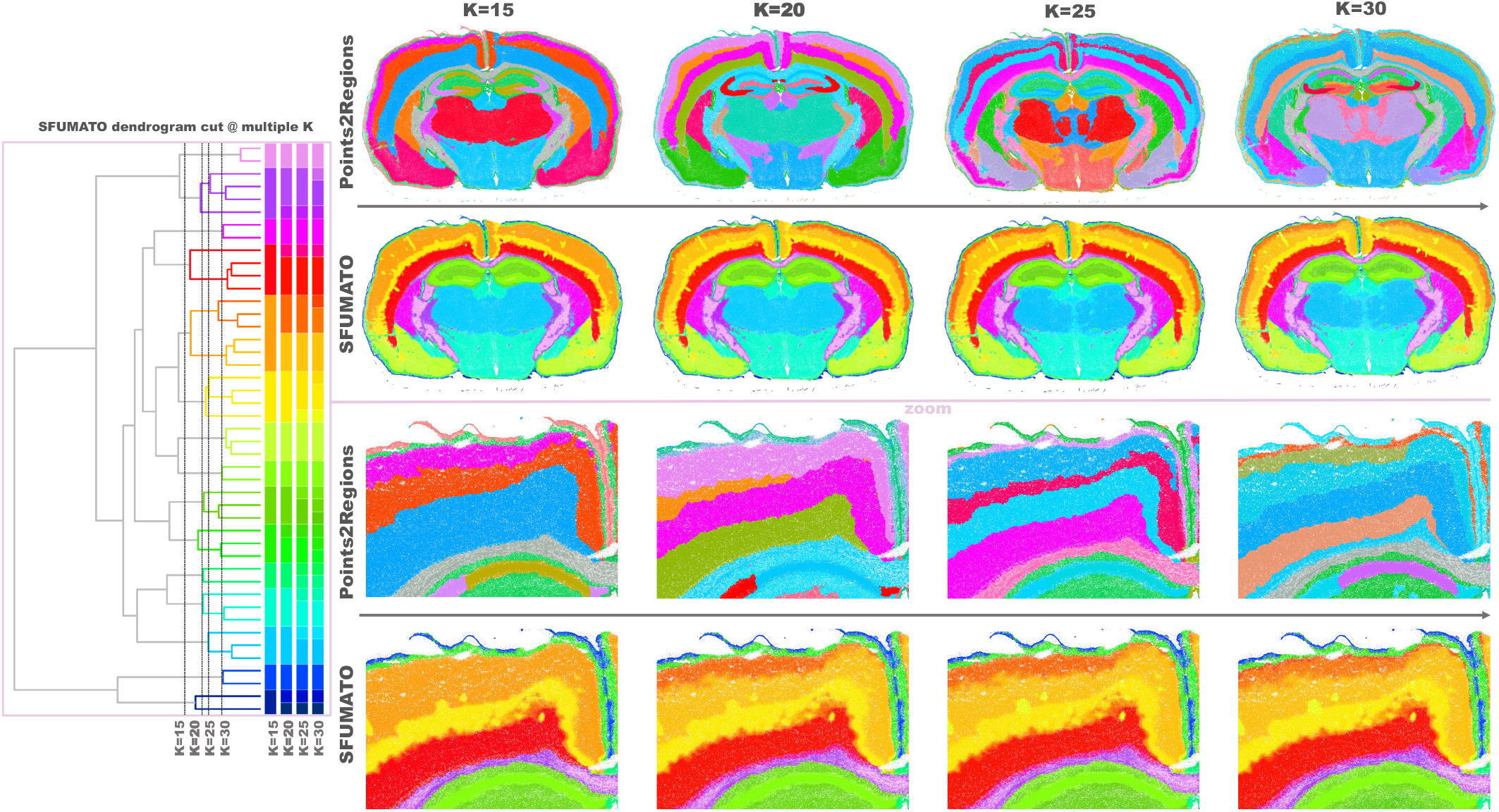
SFUMATO preserves gradual and sharp transitions across cluster resolutions. SFUMATO and Points2Regions were run on the Xenium mouse brain dataset at a niche-level bin width for *K* = 15, 20, 25, and 30 clusters. Whole-section maps show the global organisation of the mouse brain across increasing values of *K*, while cortical zoom-ins highlight local boundary behaviour. Because SFUMATO colours are inherited from a shared expression-based dendrogram, related clusters retain related colours across cuts of the hierarchy, avoiding ambiguous borders seen at clustering by Points2Regions. In transition zones, SFUMATO displays visible colour mixing between related cluster colours, whereas sharper transcriptional differences appear as stronger colour contrasts. The SFUMATO and Points2Regions niche-level results can be explored interactively for *K* = 15, 20, 25, and 30 at https://sfumato.serve.scilifelab.se/xenium_mouse_SFUMATO_niche_15.tmap, https://sfumato.serve.scilifelab.se/xenium_mouse_SFUMATO_niche_20.tmap, https://sfumato.serve.scilifelab.se/xenium_mouse_SFUMATO_niche_25.tmap, and https://sfumato.serve.scilifelab.se/xenium_mouse_SFUMATO_niche_30.tmap, respectively.

This hierarchical consistency is particularly evident in the cortical region shown in the zoomed panels of Fig. 3. In Points2Regions, the same cortical area appears with different categorical boundaries as *K* increased, making it difficult to distinguish stable anatomical or transcriptional boundaries from *K*-dependent splits. Several values of *K* therefore have to be inspected together to infer that part of this region likely corresponds to a transition zone rather than to a single sharp boundary. In SFUMATO, the same area is represented by gradual changes among related yellow-orange shades, with visible colour mixing in the transition zone. This behaviour reflects both the proximity of the corresponding clusters in the expression-based hierarchy and the probabilistic nature of the SFUMATO assignments, conveying a gradual change in transcriptional composition without imposing a visually abrupt boundary.

At the same time, SFUMATO also preserves sharper transitions where the expression profiles are more distinct. In the lower cortical region, strong hue changes from red/purple to green are spatially localised and consistently maintained across values of *K*, supporting the presence of more clearly separated transcriptional patterns. Small yellow structures, consistent with the vessel-associated patterns observed in the previous analysis, are also visible in SFUMATO but are less apparent in the Points2Regions maps. Together, these examples show that SFUMATO can display gradual transitions and sharp contrasts within the same tissue section, while maintaining a hierarchically consistent and visually robust interpretation across different choices of the number of clusters.

### 3.3 SFUMATO identifies cell types without cell segmentation

We next evaluated whether SFUMATO can identify biologically meaningful cell types without relying on cell segmentation. SFUMATO was run on the Xenium mouse brain at cell scale using the parameters reported in Supplementary Tables 2–3. To enable a direct comparison with the graph-based clustering provided by the Xenium analysis pipeline (here referred to as the Xenium reference cell typing), the shared dendrogram was cut to obtain *K* = 45 clusters, matching the number of graph-based clusters [20], resulting in the maps and zoomed-in regions shown in Fig. 4A and B. SFUMATO masks were extracted from posterior probability maps using the parameters reported in Supplementary Table 6.

**Figure 4:**
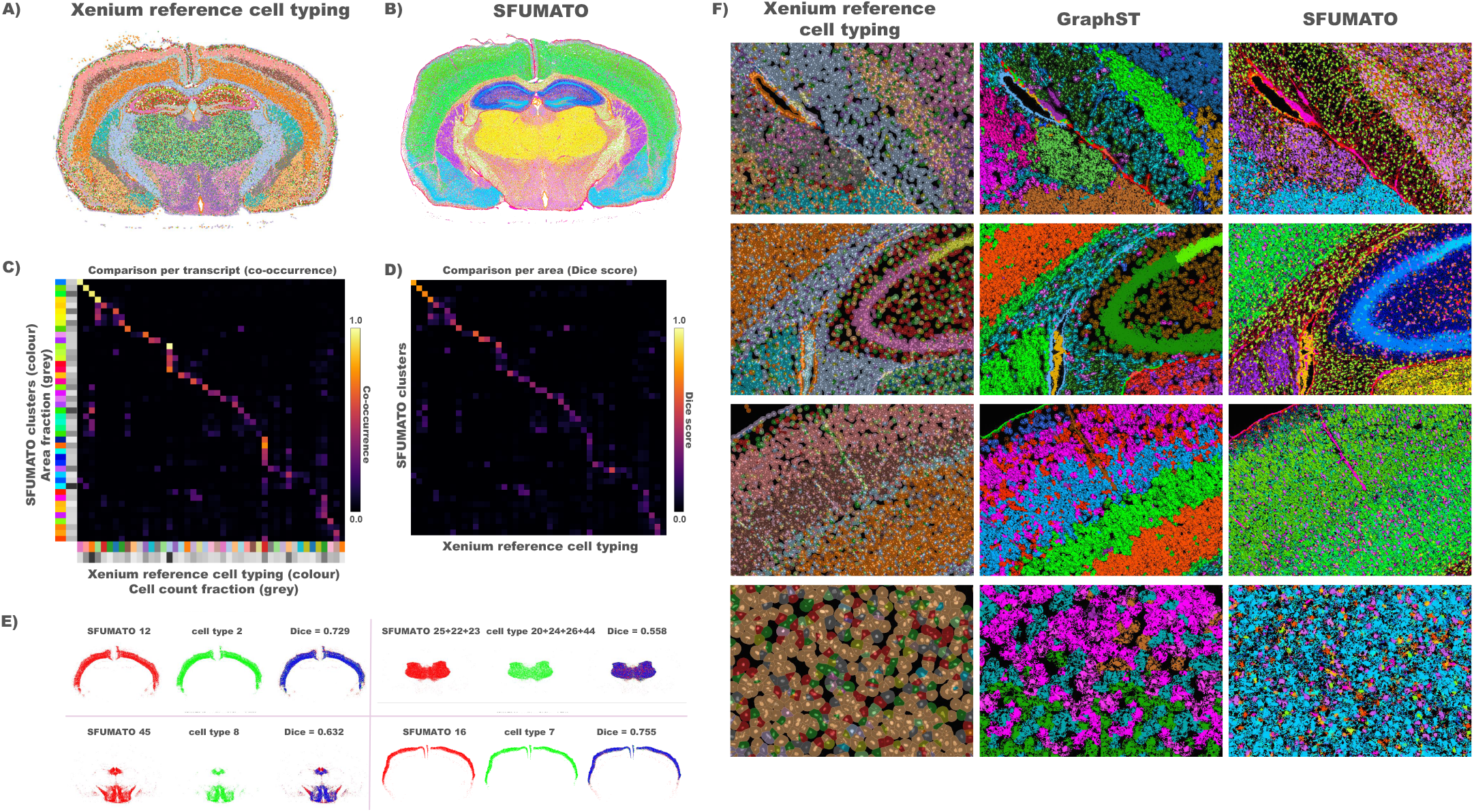
Segmentation-free identification of cell types by SFUMATO. Comparison between (**A**) the Xenium reference cell typing and (**B**) SFUMATO using *K* = 45 clusters. (**C**) The transcript-level row-normalised co-occurrence matrix exhibits a predominantly diagonal structure, indicating strong correspondence between SFUMATO clusters and Xenium reference cell types. Greyscale annotations report the relative abundance of each population, expressed as SFUMATO area fraction (rows) and Xenium reference cell type-count fraction (columns). (**D**) The corresponding Dice score matrix compares semantic segmentation masks derived from SFUMATO posterior probabilities with Xenium reference cell type masks, showing a spatial agreement consistent with the transcript-level correspondence. (**E**) Representative examples of one-to-one and grouped correspondences are shown below the matrices (SFUMATO semantic regions in red, Xenium reference cell type masks in green, and their overlap in blue). In the grouped example, transcriptionally related clusters are merged before Dice computation to account for cases in which multiple neighbouring Xenium reference cell types correspond to multiple neighbouring SFUMATO clusters. (**F**) Qualitative comparison between the Xenium reference cell typing, GraphST, and SFUMATO. The Xenium reference cell typing, SFUMATO results and GraphST results can be explored interactively at https://sfumato.serve.scilifelab.se/xenium_mouse_SFUMATO_cell.tmap and https://sfumato.serve.scilifelab.se/xenium_mouse_GRAPHST_cell.tmap, respectively.

The transcript-level co-occurrence matrix (Fig. 4C) shows a predominantly diagonal structure, indicating a high level of agreement between the graph-based Xenium reference cell types and the segmentation-free clusters identified by SFUMATO. Despite relying exclusively on transcriptomic information, SFUMATO recovered most of the major cell populations proposed by the Xenium reference cell typing.

As observed in the comparison with Points2Regions and FICTURE (Fig. 2), several many-to-one correspondences were also present when comparing SFUMATO to the Xenium reference cell typing. Importantly, these correspondences primarily involved SFUMATO clusters with similar colours, indicating that the merged Xenium reference cell types often corresponded to neighbouring clusters in the SFUMATO expression-based hierarchy rather than to transcriptionally unrelated populations.

The greyscale annotations alongside the co-occurrence matrix report the relative abundance of each population, expressed as Xenium reference cell type-count fraction and as SFUMATO area fraction for the corresponding segmentation-free clusters. Overall, agreement was highest for abundant cell populations, whereas correspondence decreased for progressively rarer populations. Nevertheless, even for low-abundance cell types, the deterioration in agreement remained limited. The least correlated region of the matrix, located in the lower-right corner, corresponded to Xenium reference cell types and SFUMATO clusters that both represented only a small fraction of the tissue.

Using the same ordering of SFUMATO clusters and Xenium reference cell types, we computed a Dice score matrix by comparing semantic segmentation masks extracted from each SFUMATO cluster with Xenium reference cell type masks (Fig. 4D). In contrast to the co-occurrence matrix, which measures transcript-level correspondence, the Dice score additionally evaluates the spatial overlap between the semantic regions recovered by the two methods.

The Dice score matrix exhibits the same predominantly diagonal structure observed in the co-occurrence matrix, confirming that the strongest transcript-level correspondences also correspond to the largest spatial overlap. The highest Dice score (0.755) was obtained between SFUMATO cluster 16 and Xenium reference cell type 7, indicating excellent agreement despite the absence of cell segmentation in the SFUMATO pipeline.

In regions exhibiting many-to-one correspondences, Dice scores were substantially lower despite relatively strong transcript-level agreement. This behaviour is expected, as the Dice coefficient assumes a one-to-one correspondence between regions. Interestingly, SFUMATO clusters 22, 23, and 25 showed correspondence with multiple Xenium reference cell types (20, 24, 26, and 44). When evaluated individually, these regions produced lower Dice scores because the corresponding spatial structures were split across several labels. After grouping SFUMATO clusters 22, 23, and 25 and Xenium reference cell types 20, 24, 26, and 44 according to the correspondences identified in the transcript-level comparison, the resulting Dice score was 0.558. This suggests that SFUMATO and the Xenium reference cell typing produce different subdivisions of the same transcriptomic region.

Supplementary Figure 7 shows the grouped Dice evaluation, and Supplementary Figure 8 shows representative TissUUmaps screenshots of the grouped SFUMATO clusters and Xenium reference cell type regions.

Representative examples of one-to-one and grouped many-to-one correspondences are shown in Fig. 4E. These examples illustrate how grouping transcriptionally related SFUMATO clusters recovers substantially larger spatial overlap with the corresponding Xenium cell type.

Finally, Fig. 4F compares the cell-type organisation recovered by SFUMATO with the Xenium reference cell typing and with GraphST, both of which rely on prior cell segmentation. Although all three methods recover the major cellular organisation of the tissue, the comparison highlights the limitations inherent to segmentation-based approaches.

The hierarchical colour assignment further facilitates the interpretation of local transcriptional organisation. Sharp changes in gene expression are represented by abrupt colour changes, whereas gradual transcriptional transitions appear as smooth colour gradients, for example in the green transition visible in the third row of Fig. 4F. In contrast, the categorical colour assignment used by GraphST and the Xenium reference cell typing tends to emphasise discrete boundaries, making gradual changes in transcriptional composition difficult to perceive.

The last row of Fig. 4F illustrates another consequence of the segmentation-free cell-typing. Regions lacking a detectable 4’,6-diamidino-2-phenylindole (DAPI) signal contain few or no segmented cells and therefore remain largely unlabelled by the Xenium reference cell typing and GraphST. Because SFUMATO operates directly on transcriptomic information rather than on segmented nuclei, it continues to recover coherent transcriptional patterns even in these regions.

### 3.4 SFUMATO can delineate pathology-associated breast cancer niches

We next evaluated whether SFUMATO could be used to identify and segment pathology-associated tissue regions from the spatial organisation of Xenium transcripts. We used the human breast cancer Xenium dataset from Janesick et al. [21], particularly Sample 1 Replicate 1, for which pathology-derived annotations delineate invasive carcinoma and two ductal carcinoma in situ regions (DCIS), denoted DCIS #1 and DCIS #2. DCIS refers to a non-invasive epithelial lesion confined within the ductal system. In this dataset, DCIS #1 and DCIS #2 correspond to distinct annotated regions selected with histology/pathology and whole-transcriptome single-cell sequencing of FFPE tissue (scFFPE-seq) guidance, while the invasive annotation marks carcinoma infiltrating the surrounding stromal tissue [21].

SFUMATO was run at niche level using the parameters reported in Supplementary Tables 2–3. We evaluated several values of *K* and inspected the corresponding expression-based dendrogram to select a single resolution that preserved the pathology-associated structures of interest. Based on this analysis, we selected *K* = 13 for semantic segmentation (Fig. 5). At this resolution, SFUMATO clusters 4 and 5 were associated with the invasive region, whereas clusters 0, 1, 2, and 3 were associated with the DCIS-annotated regions and were therefore grouped into a broader DCIS-like semantic compartment. The presence of multiple transcriptomically distinct SFUMATO clusters within the same pathological compartment suggests that a single pathological region may encompass heterogeneous transcriptional states.

**Figure 5:**
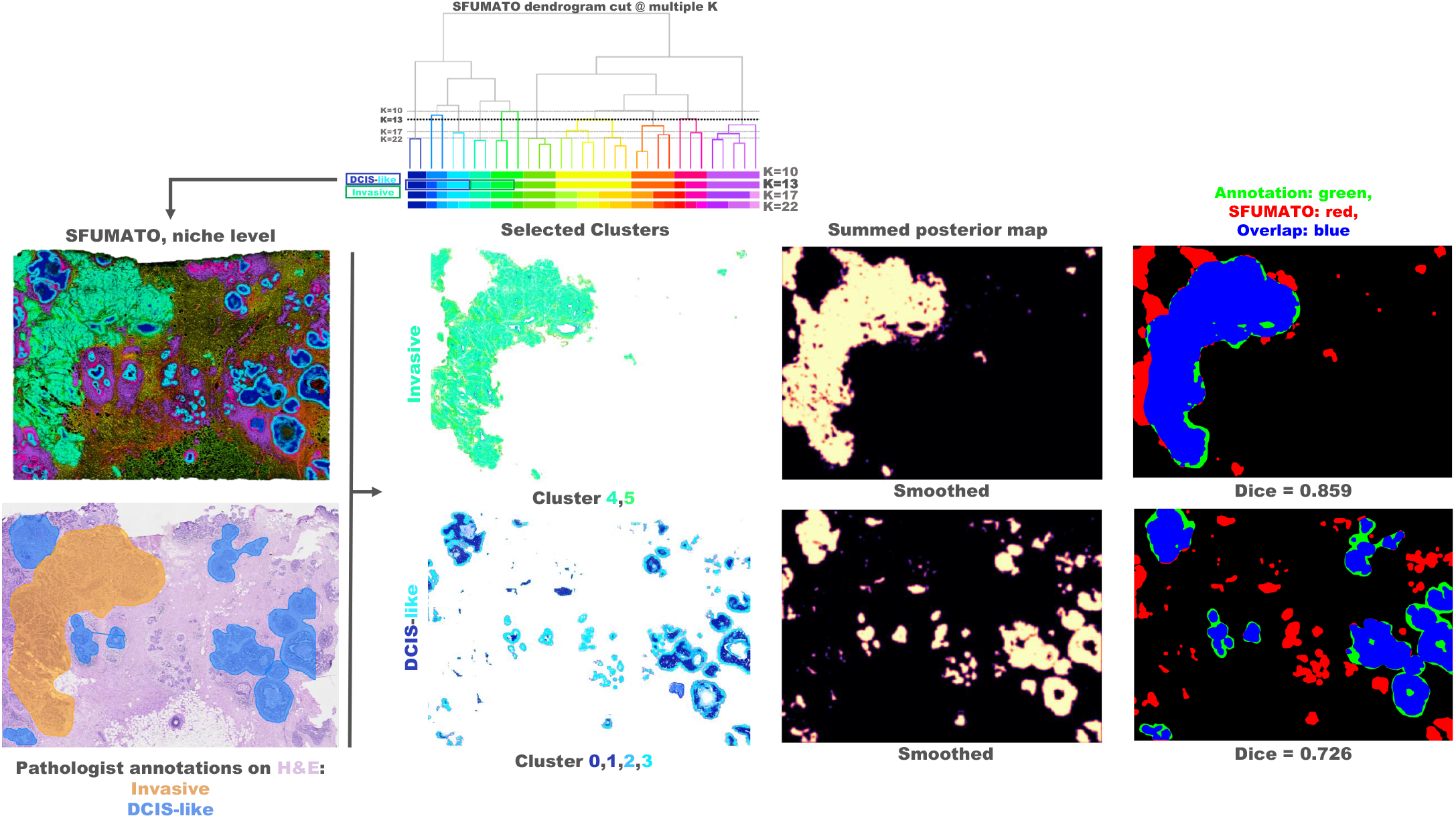
Semantic segmentation of pathology-associated breast cancer niches. SFUMATO was applied at niche level to a human breast cancer Xenium dataset and evaluated against pathology-derived annotations of invasive carcinoma and DCIS regions. Based on the expression-based dendrogram, a single resolution of *K* = 13 was selected for semantic segmentation. SFUMATO clusters 4 and 5 were grouped as an invasive-associated semantic compartment, whereas clusters 0, 1, 2, and 3 were grouped as a broader DCIS-like compartment. For each compartment, posterior probabilities were summed across the selected clusters, smoothed, and converted into binary masks by morphological reconstruction, followed by hole filling and small-object removal. Spatial concordance with the pathology annotations was quantified using the Dice similarity coefficient. In the mask overlays, regions detected only by SFUMATO are shown in red, regions present only in the pathology annotation are shown in green, and overlapping regions are shown in blue. The SFUMATO niche-level results and pathology-derived annotations can be explored interactively at https://sfumato.serve.scilifelab.se/breast_samp1_rep1_dapi_he_nichetmp.tmap.

Semantic masks were obtained from the SFUMATO posterior probability maps using the same segmentation strategy described above, with mask extraction parameters reported in Supplementary Table 6. Because the invasive and DCIS-like compartments were defined as groups of SFUMATO clusters, posterior probabilities were first summed across the selected clusters to obtain one aggregate posterior probability map per semantic compartment. The aggregate posterior probabilities were then projected into image space, smoothed with a Gaussian filter, and converted into binary masks by morphological reconstruction. Hole filling and small-object removal were then applied using spatial scales comparable to the bin size, in order to suppress isolated artefacts while preserving coherent tissue regions.

The resulting SFUMATO-derived masks were compared with the pathology annotations using the Dice similarity coefficient. The invasive compartment showed high spatial concordance with the manual annotation, with a Dice score of 0.859. The DCIS-like compartment showed lower but still substantial spatial overlap with the combined DCIS annotation, with a Dice score of 0.726. In Fig. 5, regions detected only by SFUMATO are shown in red, regions present only in the pathology annotation are shown in green, and overlapping regions are shown in blue.

Qualitatively, the invasive semantic mask closely followed the pathology-annotated invasive region, whereas the DCIS-like mask was more spatially fragmented and extended beyond parts of the manual DCIS annotations. This partial mismatch suggests that the DCIS-like SFUMATO compartment captures a broader transcript-defined pattern than the manual DCIS annotations, which may reflect heterogeneity within the annotated DCIS regions, related transcriptional programs in neighbouring structures, or differences in the spatial extent captured by transcript-based and pathology-based annotations. Together, these results support the use of SFUMATO to derive pathology-concordant semantic masks from transcript organisation alone, particularly for invasive carcinoma, while finer separation of DCIS subregions was not recovered as a stable niche-level segmentation in this setting.

### 3.5 Cell-level interpretation of breast cancer compartments

We further analysed the same breast cancer dataset at cell scale, with parameters reported in Supplementary Tables 2–3. In this analysis, we asked whether SFUMATO components could be related to broad cellular compartments described by the multimodal analysis of Janesick et al. [21]. As reference annotations, we used the cell-type labels provided in the study: Xenium transcripts were assigned to cells using nucleus-based segmentation, obtained by expanding DAPI-detected nuclei until either a maximum distance of 15 *µ*m or the boundary of a neighbouring cell was reached. The authors first identified cell types using whole-transcriptome scFFPE-seq data and then used these annotated cell profiles to assign a cell type to each segmented Xenium cell based on its expression of the 313-gene Xenium panel. For the present analysis, these detailed labels were consolidated, where appropriate, into broader biological compartments, including macrophages, myoepithelial cells, invasive tumour cells, endothelial cells, and DCIS-associated cells. The complete mapping is reported in Supplementary Table 8, and the co-occurrence matrix that aided the choice is shown in Supplementary Figure 9.

Figure 6 shows the SFUMATO dendrogram cut at *K* = 17, together with the corresponding dendrogram for colour assignment and selected cell-associated components. Fig. 6 A and B show the haematoxylin and eosin (H&E) images of the two regions highlighted in red and yellow, respectively, in the full-tissue SFUMATO map (Fig. 6 C). Panels D and E compare the Janesick-derived compartments with the corresponding SFUMATO components within these regions. The Janesick-derived compartments were coloured to match the corresponding SFUMATO components, facilitating visual comparison between the transcript-defined SFUMATO map and the segmentation-derived reference labels. The figure illustrates that broad cellular compartments can correspond to one or multiple related SFUMATO components, as reflected by their positions in the expression-based hierarchy and their related colour assignments.

**Figure 6:**
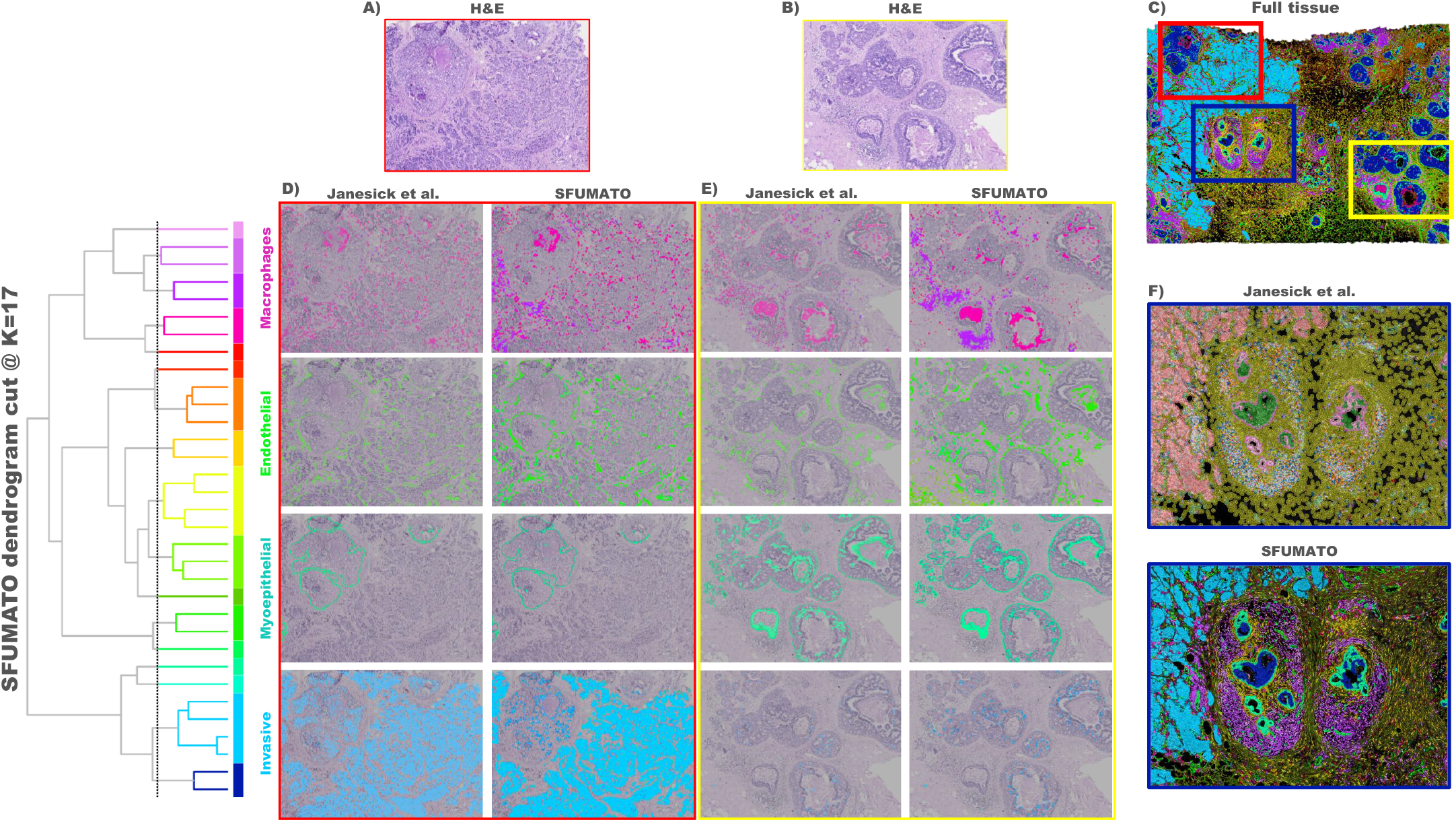
Cell-level interpretation of SFUMATO components in a breast cancer Xenium dataset. SFUMATO was applied at cell-associated scale and the resulting components were compared with Janesick-derived cell labels mapped to the Xenium cell segmentation. The dendrogram shows the expression-based hierarchy at *K* = 17 and the corresponding colour assignment. Selected SFUMATO clusters were manually related to broad cellular compartments using co-occurrence with the Janesick-derived labels and visual inspection of the spatial maps. Invasive tumour was associated with cluster 1, endothelial cells with clusters 4 and 5, myoepithelial cells with cluster 3, and macrophages with clusters 12 and 14. **(A**,**B)** H&E images of the regions highlighted by the red and yellow boxes, respectively, in the full-tissue SFUMATO map. **(C)** Full-tissue SFUMATO map showing the locations of the regions examined in the subsequent panels. **(D**,**E)** Comparison of the Janesick-derived compartments and corresponding SFUMATO components in the red- and yellow-boxed regions, respectively. **(F)** Broader comparison of the Janesick-derived cell labels and SFUMATO cell-level map in the blue-boxed region, illustrating both spatially coherent broad compartments and regions where transcript-defined SFUMATO components do not correspond one-to-one to segmentation-derived cell labels. The cell-scale SFUMATO results and Janesick-derived cell labels can be explored interactively at https://sfumato.serve.scilifelab.se/breast_samp1_rep1_dapi_he_celltmp.tmap.

Invasive tumour and myoepithelial cells showed more compact associations, mainly corresponding to SFUMATO clusters 1 and 3, respectively, and were spatially coherent with the corresponding Janesick-derived labels in the inspected regions (panels D and E in Fig. 6). Endothelial cells and macrophages were associated with multiple related SFUMATO components and showed a less strictly one-to-one correspondence. DCIS-associated cells were mainly associated with SFUMATO cluster 0, but DCIS #1 and DCIS #2 were not recovered as separate SFUMATO components, consistent with the niche-level analysis. This supports the interpretation that the DCIS subdivision reported by Janesick et al. is not directly recoverable by unsupervised analysis of the targeted Xenium transcript panel alone.

Panel F in Fig. 6 provides a broader comparison between the SFUMATO cell-level map and the Janesick-derived cell labels mapped to the Xenium segmentation. Overall, the spatial organisation of the selected broad compartments is similar between the two representations. Visual correspondence is reduced in the purple SFUMATO region, where the Janesick-derived annotation contains a dense mixture of several cell labels rather than a single dominant compartment.

In the stromal regions, differences between the two representations emerge, reflecting the different spatial units used by the two approaches: SFUMATO identifies transcript-defined domains from spatial bins, whereas the Janesick-derived reference is assigned to segmented cells. Therefore, particularly in sparse stromal areas, the two representations are not expected to have a strict one-to-one spatial correspondence.

## 4 Discussion

### A hierarchical probabilistic representation of spatial transcriptomics

SFUMATO is not simply another clustering algorithm, nor is its contribution limited to replacing a conventional clustering method with a BGM. Instead, it proposes a holistic probabilistic representation of spatial transcriptomics in which clustering, visualisation, and semantic segmentation are all derived from the same probabilistic model. Rather than treating visualisation as a post-processing step, SFUMATO makes it an integral part of the method itself.

The choice of a generative Bayesian model reflects the view that the observed transcriptomic signal arises from underlying biological organisation that should be inferred rather than imposed. Consequently, the probabilistic representation is not only used to assign cluster memberships, but also to organise the hierarchical relationships between clusters, to derive semantic regions, and to communicate uncertainty directly through visualisation.

Within this framework, once the spatial scale has been defined during feature extraction – at either the niche or cell scale – changing the number of clusters should not be interpreted as replacing one clustering with another, but rather as progressively refining the same hierarchical representation. Different values of *K* therefore correspond to different levels of biological detail rather than fundamentally different interpretations of the data. This perspective naturally motivates the use of a shared overclustered model and a common dendrogram, from which all clustering solutions are derived.

The hierarchical colour assignment further extends this idea by encoding the relationships between clusters directly into the visual representation. Similar colours indicate neighbouring positions in the expression-based hierarchy, while contrasting colours correspond to more distant branches. The subsequent probabilistic colour mixing complements this representation by revealing uncertainty in cluster assignment through gradual colour transitions, allowing biological gradients and mixed transcriptional states to emerge naturally rather than being replaced by artificial categorical boundaries.

Taken together, SFUMATO promotes the view that faithful and intuitive visualisation is not merely a means of displaying the output of an analysis, but an integral part of biological interpretation. Biological organisation is not solely composed of discrete entities separated by sharp boundaries, but also of gradual transitions and continuous variation. The name *SFUMATO* reflects this central idea: rather than suppressing biological gradients through hard assignments, the framework aims to preserve and communicate them through an interpretable probabilistic representation.

### Segmentation-free identification of cellular organisation

A fundamental characteristic of SFUMATO is that it operates directly on the spatial transcriptomic signal without requiring prior cell segmentation or underlying DAPI images. Consequently, the probabilistic model is not affected by segmentation errors, such as missing nuclei, merged or overlapping cells, or transcripts originating from cells whose nuclei fall outside the imaging plane.

Xenium graph-based clustering and GraphST rely on prior cell segmentation, and inevitably propagate segmentation errors to the downstream clustering. SFUMATO’s segmentation-free formulation enables operation directly on transcriptomic information; the observer can exploit the full transcriptomic signal, independently of image-derived cell boundaries.

In regions where neighbouring cells are fused into a single segmented object, segmentation-based methods inevitably assign the transcriptomic signal originating from different cells to a single instance, and therefore to a single cluster. In contrast, SFUMATO naturally models these situations through its probabilistic formulation: transcripts originating from spatial bins containing mixed signals contribute to multiple cluster assignments, and the resulting uncertainty is communicated through probabilistic colour mixing. Rather than forcing a hard assignment, the method explicitly represents ambiguous transcriptional states through gradual colour transitions, providing a more faithful description of complex cellular microenvironments.

In regions lacking detectable DAPI signal, SFUMATO continues to recover coherent transcriptional patterns from transcriptomic information alone: because gene expression occurs throughout the cell rather than exclusively near or within the nucleus, transcripts can be detected outside the segmented DAPI signal and form meaningful patterns.

### Area fractions as a quantitative descriptor of tissue composition

According to the Delesse principle, measurements of area fraction obtained from two-dimensional sections provide an unbiased estimator of the corresponding volume fractions (under the assumptions of the principle) [13]. Although this may require corrections in biological tissues [14], area fraction constitutes the natural quantity produced by the probabilistic formulation of SFUMATO. Cell counts and area fractions describe different biological properties and therefore should not be expected to coincide. Which quantity is more appropriate ultimately depends on the biological question being addressed. However, when the objective is to characterise the spatial occupancy of transcriptionally defined populations within a tissue, area fractions may provide a more natural descriptor than cell counts, while avoiding the dependence on image-based cell segmentation. Cell quantification from area fractions, when normalised for variation in volume of different cell types, is also more likely to correlate with cell counts derived from single cell sequencing.

Area fraction quantification arises naturally within the SFUMATO framework. Because each cluster is represented by a posterior probability map, semantic regions and their corresponding area fractions can be obtained directly from the probabilistic transcriptomics representation, without requiring cell segmentation. The probabilistic formulation therefore provides not only an uncertainty-aware clustering, but also a quantitative description of the spatial extent of transcriptional populations.

The distinction between cell counts and area fraction also provides an alternative interpretation of part of the disagreement observed between SFUMATO and the Xenium 10x Genomics cell-type annotations. While the Xenium 10x Genomics workflow describes tissue organisation through segmented cells, SFUMATO identifies transcriptomic domains directly from the spatial gene-expression signal. Consequently, discrepancies in Dice score or one-to-one cluster correspondence should not necessarily be interpreted solely as clustering errors, but may also reflect the fact that the two methods quantify different aspects of the same underlying biological organisation. In particular, transition zones and regions containing mixed transcriptional signals are naturally represented by the probabilistic formulation of SFUMATO, whereas segmentation-based approaches are required to assign these regions to discrete cellular objects.

### Semantic regions as a bridge between transcriptomics and pathology

The semantic segmentation of pathology-associated niches highlights the role of SFUMATO as an intermediate representation between spatial transcriptomics and pathology. By converting cluster-specific posterior probability maps into semantic masks, SFUMATO provides transcript-derived regions that can be directly compared with pathology annotations. In this setting, the invasive carcinoma-associated compartment showed strong spatial concordance with the manual annotation, whereas the DCIS-like compartment showed only partial overlap and appeared as a broader, more heterogeneous transcript-defined region.

This difference is informative. The DCIS #1 and DCIS #2 labels reported by Janesick et al. were obtained through an integrative analysis involving whole-transcriptome scFFPE-seq, Visium, histopathology, marker interpretation, and subsequent label transfer to Xenium, rather than through unsupervised analysis of the targeted Xenium transcript panel alone [21]. Therefore, the absence of a stable SFUMATO separation between DCIS #1 and DCIS #2 should not be interpreted simply as a segmentation or clustering failure. Instead, it suggests that some pathology-associated subregions may depend on information that is only partially captured by the targeted transcript panel.

For the same reason, the cell-level comparison on the breast cancer dataset should be used to interpret SFUMATO components biologically, rather than to evaluate SFUMATO as a direct cell-type classifier. The Janesick-derived cell labels used as reference were produced within a multimodal and segmentation-based annotation framework, whereas SFUMATO infers transcript-defined components directly from spatial bins. Dataset-specific multimodal downstream analyses can therefore complement SFUMATO when finer cell-state or cell-type interpretation is required.

### Dimensionality reduction for probabilistic clustering

Dimensionality reduction is a key preprocessing step for imaging-based spatial transcriptomics. In the Xenium analysis workflow described by Salas et al. [10], the number of retained principal components was chosen to explain over 80% of the total variance. Our experiments suggest that this criterion is not necessarily optimal when the downstream task is probabilistic clustering. Empirically, relatively compact latent representations consistently yielded more stable cluster assignments and more interpretable biological patterns. In the context of SFUMATO, dimensionality reduction is therefore not intended to maximise variance preservation or faithfully reconstruct the original feature space, but rather to produce a compact, denoised latent representation suitable for subsequent probabilistic modelling.

For this reason, the number of retained latent dimensions, *d*, should be regarded as a tunable hyperparameter rather than as a quantity determined solely by explained variance. Its optimal value depends on the characteristics of the dataset, the biological question under investigation, and the desired level of transcriptional detail.

We adopted TruncatedSVD rather than conventional PCA for feature reduction because it operates directly on sparse matrices without requiring matrix densification, making it particularly suitable for highly multiplexed spatial transcriptomics data. Although TruncatedSVD differs mathematically from conventional PCA, as it does not centre the data before computing the singular value decomposition, the log-transformed and row-normalised feature representation produced by the Points2Regions feature extraction substantially reduces the practical impact of this difference. TruncatedSVD therefore retains computational advantages for sparse matrices while providing a latent representation comparable to that obtained by PCA in the present setting.

We also explored alternative dimensionality reduction strategies, including nonlinear embedding methods and joint probabilistic formulations inspired by BFAST [5], in which latent representation learning and clustering are performed simultaneously. While conceptually attractive, these approaches substantially increased computational complexity without producing commensurate improvements in clustering quality. In particular, the computational efficiency of SFUMATO relies on the use of diagonal covariance matrices within the BGM, allowing scalable inference on large spatial transcriptomics datasets. We therefore found the sequential TruncatedSVD–BGM pipeline to provide the most favourable compromise.

### Limitations

The hierarchical colour assignment adopted by SFUMATO inevitably provides a one-dimensional representation of a complex hierarchical organisation. The optimally ordered dendrogram ensures that neighbouring colours correspond to neighbouring clusters in the expression-based hierarchy, and different shades of similar colours indicate transcriptionally related populations. However, colour should not be interpreted as a quantitative measure of transcriptional similarity, but rather as an intuitive representation of the hierarchical relationships between clusters.

Furthermore, the number of perceptually distinguishable colours is inherently limited. As the number of clusters increases, maintaining sufficient visual contrast between all clusters becomes progressively more difficult, despite the increased stability and hierarchical consistency provided by the shared colour assignment.

A second limitation concerns parameter selection. The spatial bin width, the smoothing factor used during feature extraction, and the dimensionality of the latent representation all influence the resulting clustering and should be tuned according to the biological question of interest. Similarly, the Gaussian smoothing and morphological reconstruction thresholds used to extract semantic regions from the posterior probability maps remain user-defined parameters that may require dataset-specific optimisation.

Finally, although posterior probabilities are retained throughout the SFUMATO pipeline, the present work exploits them primarily for uncertainty-aware visualisation, semantic segmentation, and area estimation. Their potential for downstream statistical analyses remains largely unexplored.

### Future perspectives

Several directions could further extend the capabilities of SFUMATO. First, the posterior probabilities inferred by the BGM contain substantially richer information than is currently exploited. Beyond uncertainty-aware visualisation, semantic segmentation, and area estimation, posterior probabilities could be incorporated directly into downstream analyses, for example by weighting differential gene expression analyses according to cluster-assignment confidence, rather than relying exclusively on hard labels. More broadly, these results support the use of SFUMATO as a transcript-derived layer in multimodal pathology workflows. In particular, the continuous posterior probability maps provide a compact representation of highly multiplexed spatial transcriptomic information, which could be combined with histology-derived features, whole-transcriptome references, or externally derived cell-state labels in future multimodal extensions.

From a methodological perspective, the probabilistic clustering and visualisation framework of SFUMATO is independent of the feature extraction stage. Although the present work relies on a Points2Regions-inspired representation, the framework could be readily combined with alternative feature representations, including convolutional neural network or foundation-model embeddings, or lower-resolution spatial transcriptomics. This flexibility makes SFUMATO applicable beyond the specific feature extraction strategy considered here.

Finally, the hierarchical colour assignment naturally lends itself to interactive visualisation tools. We envision a dedicated TissUUmaps [11, 12] plug-in module capable of dynamically adapting colour contrast to user-selected regions of interest, thereby improving the visualisation of local transcriptional patterns while preserving the global hierarchical consistency of the colour assignment.

## Acknowledgements

This work was financed through a project grant from the Swedish Research Council (2024-05150) to CW.

The computations were enabled by resources in project NAISS2026-4-793 provided by the National Academic Infrastructure for Supercomputing in Sweden (NAISS) at Alvis, funded by the Swedish Research Council through grant agreement No. 2022-06725.

## Data Availability

The SFUMATO analysis pipeline, together with example notebooks that reproduce the analyses presented in this study, is publicly available at https://github.com/ale2sia-g/SFUMATO. Interactive visualizations of the results are available through TissUUmaps at https://sfumato.serve.scilifelab.se/.

The datasets analysed in this study are publicly available on the website https://www.10xgenomics.com/datasets, as described in the Methods section.

## Materials and correspondence

Correspondence and requests for materials should be addressed to CW.

## Supplementary Material

### Datasets and reference annotations

**Table 1:** Summary of Xenium datasets used in this study. The table reports the public Xenium datasets used for SFUMATO evaluation, the sample analysed, the available imaging/reference information, and the role of each dataset in the study. Transcript counts refer to the number of rows in the transcript coordinate files used as input to SFUMATO.

| Dataset | Sample analysed | Sample type | Panel / input size | Available reference information | Use in this study |
| --- | --- | --- | --- | --- | --- |
| Xenium mouse brain | Full coronal section | Fresh frozen mouse brain | Xenium Mouse Brain Gene Expression Panel; 248 genes; 42,196,647 transcript records used as SFUMATO input | Xenium morphology image, cell segmentation, cell-feature matrix, and Xenium Onboard Analysis outputs, including UMAP, clustering, and differential expression results | Evaluation of SFUMATO colour encoding, hierarchical consistency across values of $K$, and segmentation-free recovery of cell-associated spatial organisation |
| Human breast cancer Xenium dataset | In situ Sample 1, Replicate 1 | FFPE human breast cancer section | Xenium Human Breast Gene Expression Panel; 280 genes plus 33 custom genes, for a total of 313 targeted genes; 42,433,535 transcript records used as SFUMATO input | Xenium morphology/cell segmentation outputs, post-Xenium H&E imaging, pathology-derived annotations, and Janesick-derived cell labels from the multimodal analysis | Evaluation of pathology-associated semantic segmentation and cell-level interpretation of SFUMATO components in relation to Janesick-derived annotations |

These input transcript counts may differ from summary metrics reported by 10x Genomics, because the latter depend on the specific filtering, quality-control, and assignment criteria used in the Xenium output summaries.

### SFUMATO parameters

**Table 2:** Feature extraction and clustering parameters used for SFUMATO analyses. The bin width is reported in Xenium image pixels (0.2125 *µ*m/px). The smoothing factor corresponds to the Points2Regions smoothing parameter and defines the maximum smoothing width relative to the bin width. *K*_BGM_ denotes the number of components used to fit the overclustered BGM, whereas *K* values indicate the final clustering resolutions obtained by cutting the shared dendrogram.

| Run / figure | Dataset | Bin width (px) | Smoothing factor | Min genes/bin | SVD Di-mensions ($d$) | $K_{\text{BGM}}$ | $K$ values |
| --- | --- | --- | --- | --- | --- | --- | --- |
| mousexeniumcell;<br>Fig. 2, Fig. 4 | Mouse brain | 7 | 5 | 4 | 10 | 68 | 45 |
| mousexeniumniche;<br>Fig. 3 | Mouse brain | 40 | 8 | 120 | 10 | 45 | 15, 20, 25, 30 |
| breastxenium_niche;<br>Fig. 5 | Human breast cancer | 40 | 8 | 200 | 10 | 33 | 10, 13, 17, 22 |
| breastxenium_cell;<br>Fig. 6 | Human breast cancer | 10 | 6 | 20 | 10 | 33 | 13, 17, 20, 22 |

**Table 3:** Bayesian Gaussian mixture, hierarchical merging, and colour-assignment parameters used for SFUMATO analyses. The table reports parameters used after feature extraction, including BGM overclustering, hierarchical merging, and colour assignment.

| Run | BGM over-sampling | Weight prior | Merge method | Merge metric | Colormap |
| --- | --- | --- | --- | --- | --- |
| mousexeniumcell | 1.5 | 1 | Complete | Cosine distance | `gist_ncar`, range 0.08–0.92 |
| mousexeniumniche | 1.5 | 1 | Complete | Cosine distance | `gist_ncar`, range 0.08–0.92 |
| breastxenium_niche | 1.5 | 1 | Complete | Cosine distance | `gist_ncar`, range 0.08–0.92 |
| breastxenium_cell | 1.5 | 1 | Complete | Cosine distance | `gist_ncar`, range 0.08–0.92 |

### Competing method parameters

Points2Regions was run using the same feature extraction parameters as SFUMATO for each corresponding analysis, as reported in Supplementary Table 2.

The Xenium analysis workflow outputs were used as platform-derived reference annotations and were not re-estimated in this study. According to the 10x Genomics documentation, Xenium Onboard Analysis performs cell segmentation, assigns transcripts to cells, and runs secondary analysis on the resulting single-cell data, including PCA, UMAP, clustering, and differential expression. The clustering outputs include graph-based and K-means results.

### FICTURE

**Table 4:**
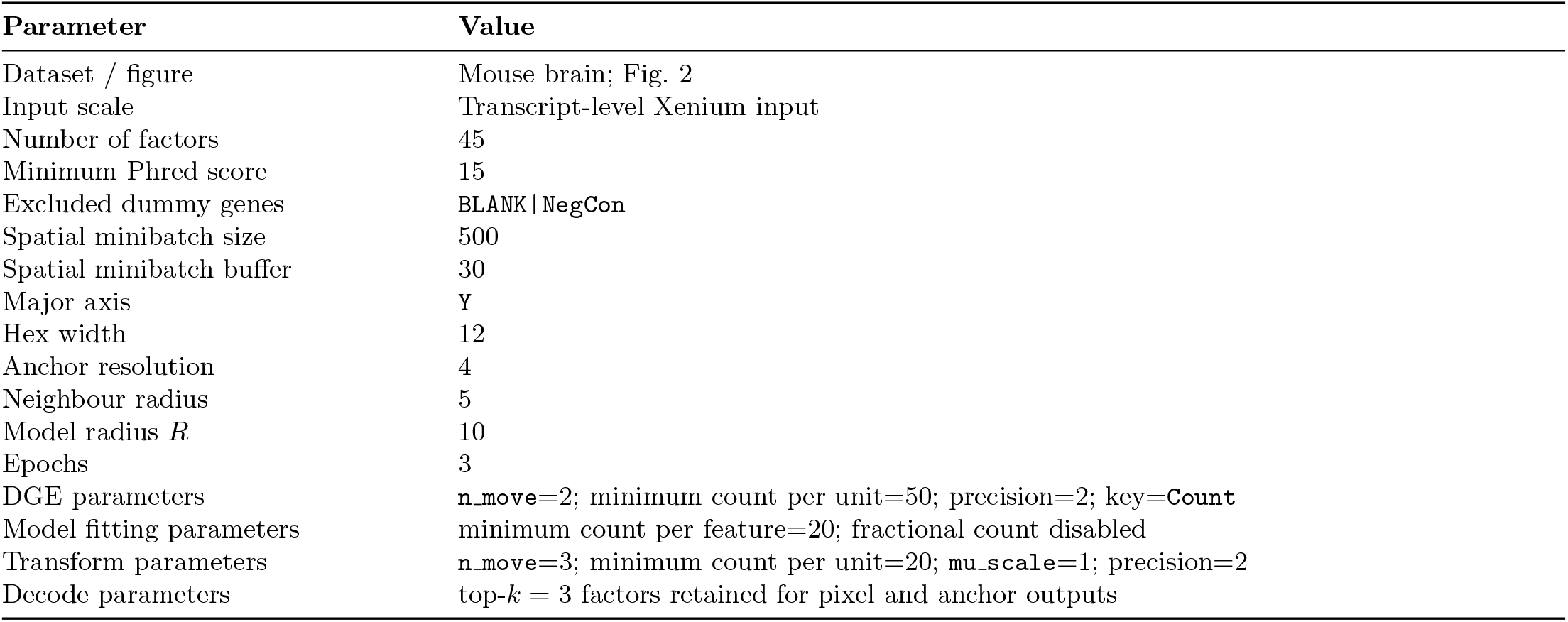
FICTURE parameters used for comparison with SFUMATO. FICTURE was run as a segmentation-free probabilistic factorization method on the Xenium mouse brain dataset.

### GraphST

**Table 5:** GraphST parameters used for comparison with SFUMATO. GraphST was applied to a cell-level representation derived from Xenium cell segmentation and clustered into 45 domains for comparison with SFUMATO and Xenium graph-based cell annotations.

| Parameter | Value |
| --- | --- |
| Dataset / figure | Mouse brain; Fig. 4 |
| Input scale | Cell-level `h5ad` representation derived from Xenium cell segmentation |
| Number of clusters | 45 |
| GraphST training | Default GraphST training procedure |
| Clustering method | Leiden |
| Resolution search | 0.01 to 5.0 |
| Resolution increment | 0.005 |
| Refinement | False |
| Output labels | `domain` clusters exported per cell |

### Semantic segmentation settings

**Table 6:**
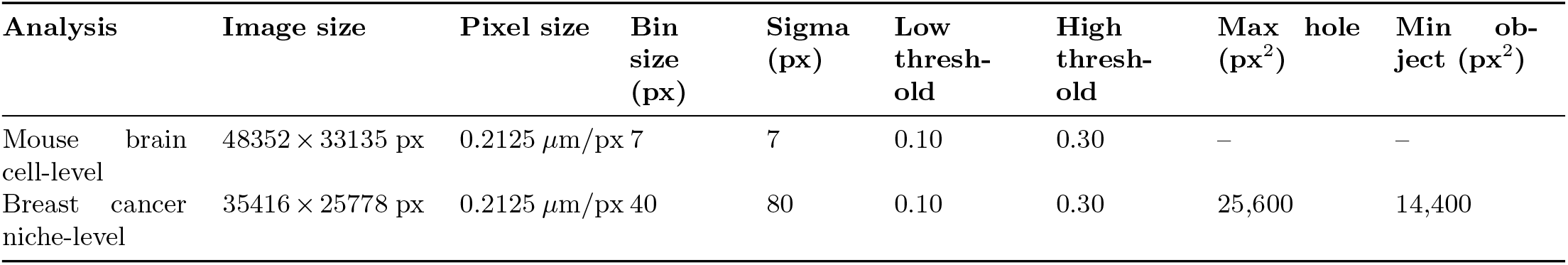
Semantic segmentation parameters used to extract binary masks from SFUMATO posterior probability maps. Posterior probability maps were projected into image space, smoothed with a Gaussian filter, and thresholded by morphological reconstruction using low and high probability thresholds. For the breast cancer niche-level analysis, holes smaller than (4 *b*)^2^ pixels and connected objects smaller than (3 *b*)^2^ pixels were removed, where *b* denotes the bin size, corresponding to 25,600 and 14,400 px^2^, respectively.

### Grouped Dice evaluation

**Figure 7:**
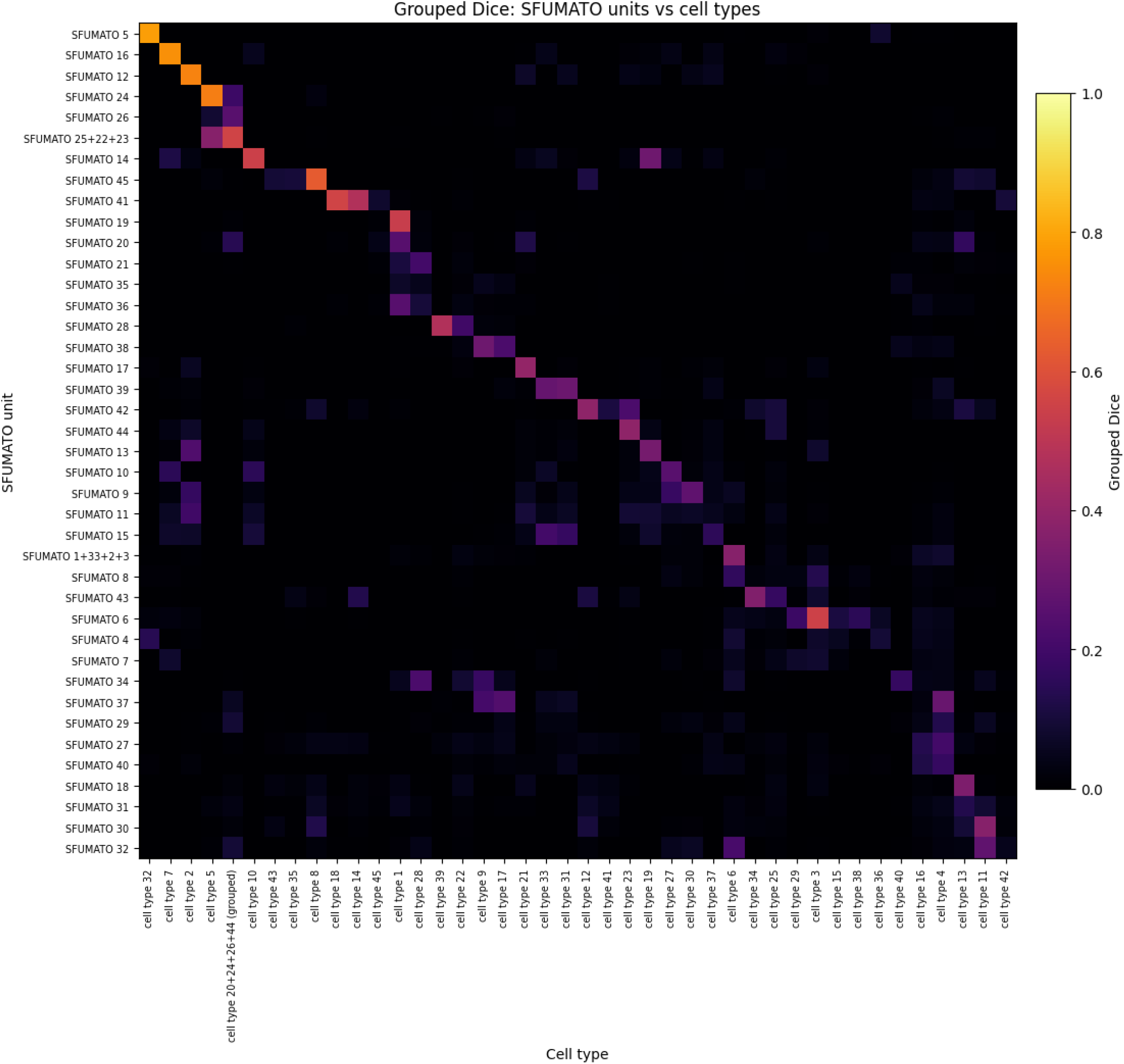
Grouped Dice comparison between SFUMATO semantic regions and Xenium reference cell type masks. Many-to-one correspondences identified from the transcript-level co-occurrence matrix were merged before Dice computation by combining the corresponding SFUMATO semantic masks. This grouped evaluation accounts for cases in which a single Xenium reference cell type corresponds to multiple neighbouring SFUMATO clusters in the expression-based hierarchy or vice versa.

**Table 7:** Grouped SFUMATO regions used for the grouped Dice evaluation. SFUMATO clusters were grouped when a Xenium reference cell type or cell type group corresponded to multiple SFUMATO clusters in the co-occurrence analysis before computing Dice similarity. For grouped Xenium targets, Dice was computed against the union of the corresponding cell-type masks.

| Grouped SFUMATO region | SFUMATO clusters | Target Xenium reference cell type(s) |
| --- | --- | --- |
| SFUMATO 1+33+2+3 | 1, 33, 2, 3 | cell type 6 |
| SFUMATO 25+22+23 | 25, 22, 23 | cell type 26, cell type 44, cell type 20, cell type 24 |

**Figure 8:**
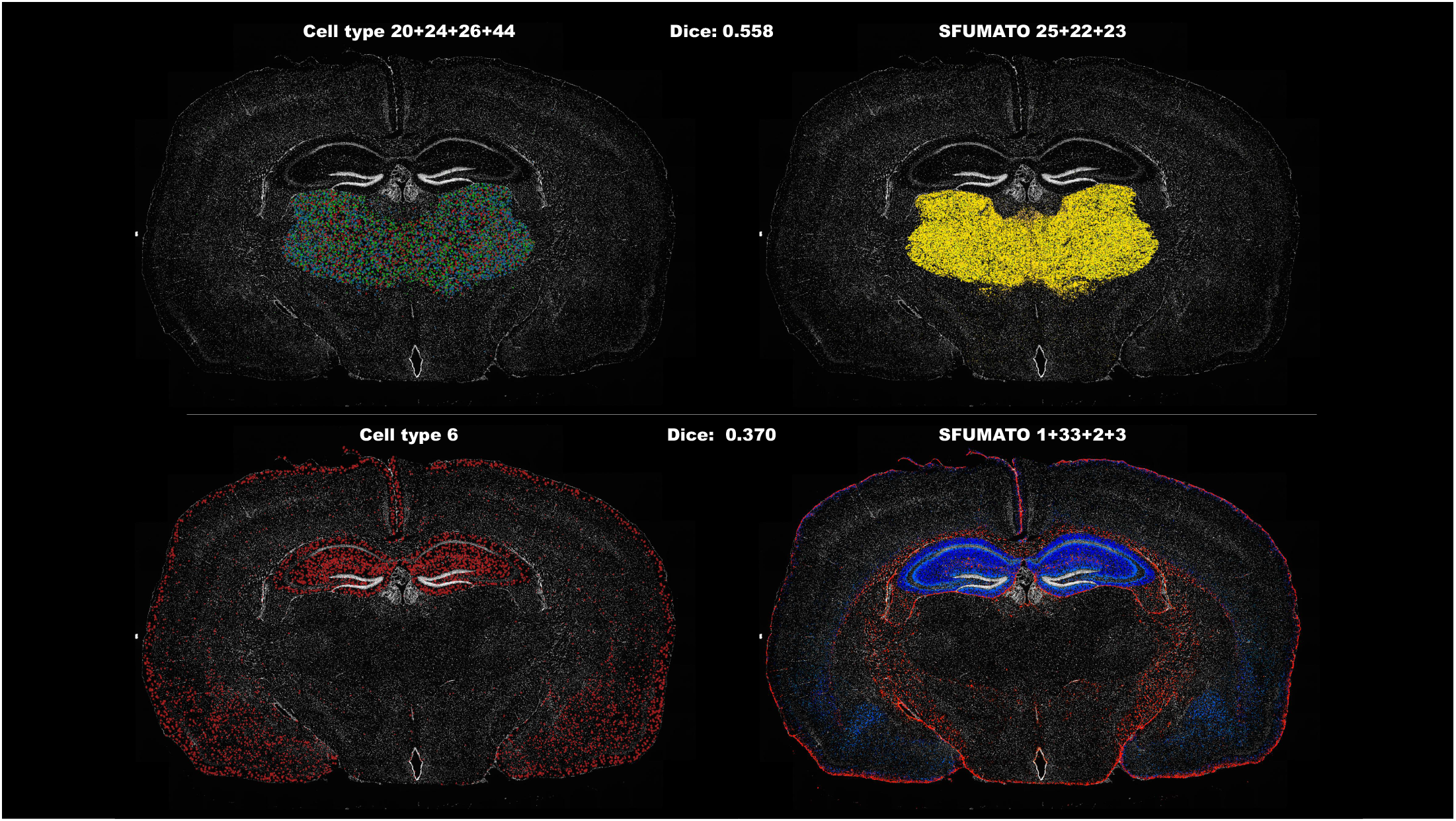
Interactive inspection of SFUMATO clusters and Xenium reference cell types in TissUUmaps. Representative regions used to inspect the spatial correspondence between SFUMATO semantic masks and Xenium reference cell type annotations. Screenshots were generated from the interactive TissUUmaps project available at https://sfumato.serve.scilifelab.se/xenium_mouse_SFUMATO_cell.tmap.

### Breast cell-level compartment mapping

**Table 8:** Broad compartment mapping used for cell-level interpretation in the breast cancer dataset. Janesick-derived labels were collapsed into broad compartments where appropriate and related manually to selected SFUMATO clusters using co-occurrence with the reference labels and visual inspection of spatial maps.

| Broad compartment | Janesick-derived labels collapsed | SFUMATO clusters |
| --- | --- | --- |
| Macrophages | Macrophages_1; Macrophages_2 | 12, 14 |
| Myoepithelial cells | Myoepi_ACTA2+; Myoepi_KRT15+ | 3 |
| Invasive tumour cells | Invasive_tumour | 1 |
| Endothelial cells | Endothelial | 4, 5 |
| DCIS-associated cells | DCIS #1; DCIS #2 | 0 |

**Figure 9:**
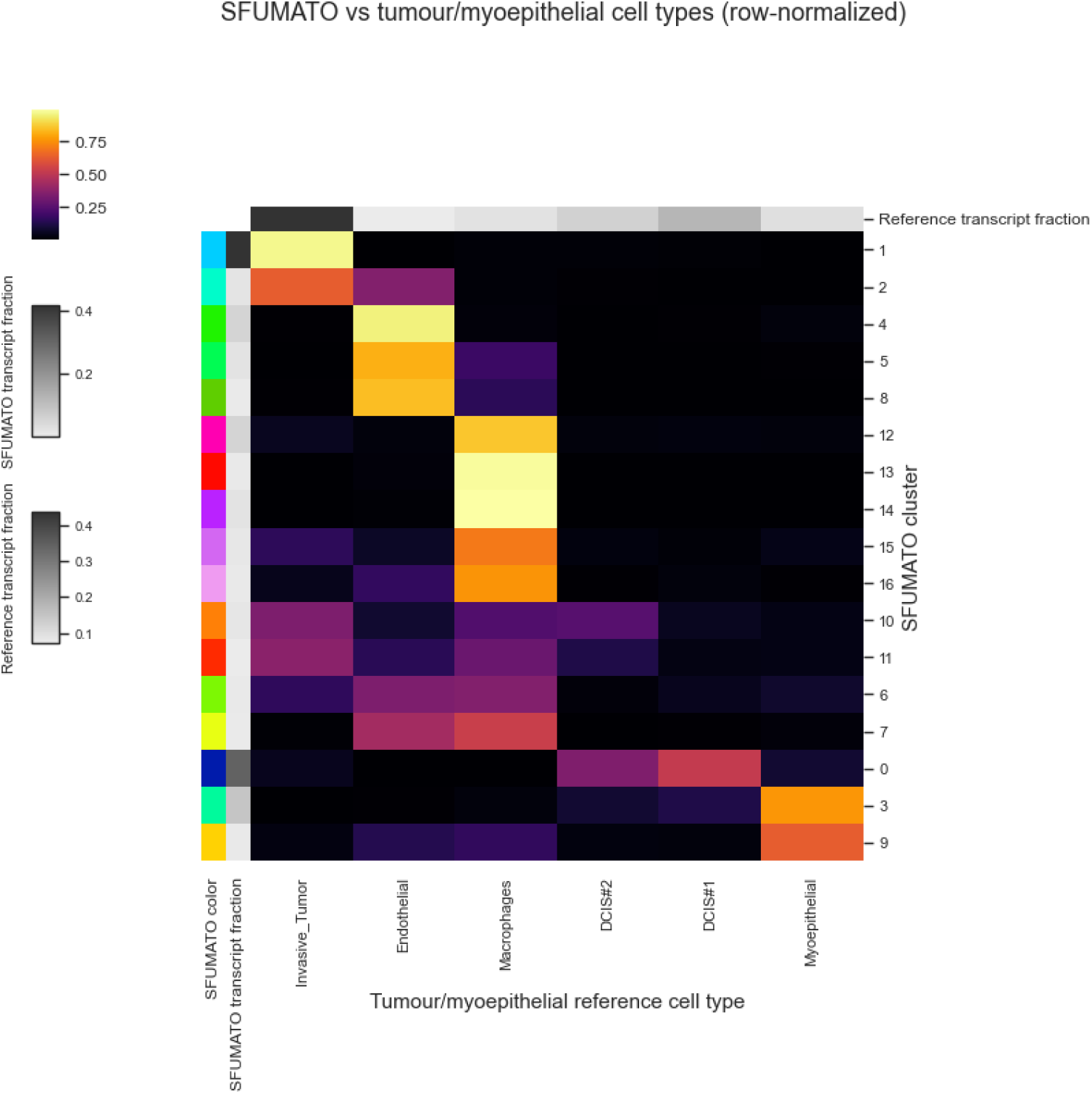
Co-occurrence between SFUMATO clusters and selected cell compartments. Detailed Janesick-derived cell-type labels were manually collapsed into broader compartments based on their biological relationships, co-occurrence patterns with SFUMATO clusters, and spatial distribution in the tissue.

